# AAVone: A Cost-Effective, Single-Plasmid Solution for Efficient AAV Production with Reduced DNA Impurities

**DOI:** 10.1101/2025.01.07.631712

**Authors:** Rongze Yang, Ngoc Tam Tran, Taylor Chen, Mengtian Cui, Yuyan Wang, Tapan Sharma, Yu Liu, Jiantao Zhang, Xinxu Yuan, Danmeng Zhang, Cuiping Chen, Zhen Shi, Leming Wang, Yuling Dai, Haniya Zaidi, Jiarui Liang, May Chen, Dabbu Jaijyan, Huan Hu, Bing Wang, Cheng Xu, Wenhui Hu, Guangping Gao, Daozhan Yu, Phillip W.L. Tai, Qizhao Wang

## Abstract

Currently, the most common approach for manufacturing GMP-grade adeno-associated virus (AAV) vectors involves transiently transfecting mammalian cells with three plasmids that carry the essential components for production. The requirement for all three plasmids to be transfected into a single cell and the necessity for high quantities of input plasmid DNA, limits AAV production efficiency, introduces variability between production batches, and increases time and labor costs. Here, we developed an all-in-one, single-plasmid AAV production system, called AAVone. In this system, the adenovirus helper genes (*E2A*, *E4orf6*, and *VA RNA*), packaging genes (*rep* and *cap*), and the vector transgene cassette are consolidated into a single compact plasmid with a 13-kb backbone. The AAVone system achieves a two- to four-fold increase in yields compared to the traditional triple-plasmid system. Furthermore, the AAVone system exhibits low batch-to-batch variation and eliminates the need for fine-tuning the ratios of the three plasmids, simplifying the production process. In terms of vector quality, AAVs generated by the AAVone system show similar *in vitro* and *in vivo* transduction efficiency, but a substantial reduction in sequences attributed to plasmid backbones and a marked reduction in non-functional snap-back genomes. In Summary, the AAVone platform is a straightforward, cost-effective, and highly consistent AAV production system – making it particularly suitable for GMP-grade AAV vectors.

## INTRODUCTION

Adeno-associated virus (AAV) vectors have become the preferred gene delivery vehicles because of their efficient transduction, long-term gene expression, low immunogenicity, and favorable safety profile. They have been successfully utilized in numerous clinical trials and have received regulatory approvals as drugs for human use.^1–3^ Although AAV production systems based on herpes simplex virus, vaccinia virus, and adenovirus (AdV) have been developed, they have not been widely adopted because of concerns related to potency, quality, safety, and processing complexity.^4–11^ The mainstay technology for AAV vector production with human cell lines (HEK293) is still the triple-plasmid (tri-plasmid) transfection method, which involves co-transfecting cells with three plasmids containing the necessary *cis* and *trans* components.^12^ The inverted terminal repeats (ITRs) are the only *cis* elements of viral origin and are required for AAV replication and packaging. Within the AAV vector plasmid that carries the transgene cassette, also called the *cis* plasmid, the gene of interest (GOI) is positioned between two ITRs. The AAV genes necessary for production, *rep* and *cap*, are supplied by a separate plasmid called the *trans* plasmid. Among the four non-structural Rep proteins encoded by the wildtype AAV (wtAAV) genome, only Rep78 and Rep52 are necessary for AAV vector production.^6^ The capsid proteins VP1, VP2, and VP3 are required for proper capsid assembly. Additionally, assembly-activating protein (*AAP*) and membrane-associated accessory protein (*MAAP*) genes, which are embedded within the *cap* gene in different open reading frames (ORFs), aid in viral capsid assembly and egress.^13–15^ To facilitate AAV vector production, a third plasmid is used to provide essential AdV genes. This “helper” plasmid expresses the *E2A*, *E4orf6*, and *VA RNA* genes.^16^ AAV vectors produced using the tri-plasmid method have demonstrated safety, convenience, and effectiveness. However, this method has limited scalability,^17,18^ and requires co-transfection of all three plasmids into a single cell.^19^ Moreover, the requirement of high quantity of plasmid DNA (pDNA) and the need for ratio optimization among the three plasmids not only increases time and labor costs, but also introduces variability between production batches.^20–22^

In addition to packaging intact GOI genomes, AAV can also encapsulate nucleic acid contaminants, including incomplete AAV vector genomes, producer plasmid genomes, and host cells DNAs (hcDNAs).^23^ The tri-plasmid systems yield a relatively high amount of impurities linked to the input plasmids. These impurities can encompass sequences related to the bacterial origin of replication and antibiotic resistance genes.^10,17,23–25^ The level of DNA impurities encapsulated in AAV vector products varies significantly, depending on the vector design and production process. Self-complementary AAVs (scAAVs) have shown that as much as 26% of the total capsids can contain backbone plasmid DNA.^26^ For standard single-stranded AAVs (ssAAVs), this figure can be as high as 10%.^23,27–29^ This concern has become more important, since recent trials have revealed significant safety concerns at doses exceeding 1E14 vg/kg.^30,31^ For instance, even if 95% of the packaged DNA within a clinical preparation were fully functional vectors, a dose of 1E14 vg/kg would comprise around 5E12 vg/kg of non-therapeutic vector.^23^ The use of an “oversized” backbone can significantly reduce residual plasmid backbone packaging.^32,33,34^

To address the increasing demand for AAV gene therapies, it is crucial to make improvements to current production systems. One approach is to develop more efficient upstream processes, including new cell lines and transfection reagents.^17,35,36^ Another approach is to optimize the design and function of the helper and *cis* plasmids used in production systems.^12,37,38^ A third approach is to streamline the process to optimize the transient-transfection step by reducing the number of plasmids required. In a dual-plasmid system, one plasmid (pDG) carries the necessary AdV helper genes, as well as the AAV *rep* and *cap* genes, while the other contains the GOI cassette.^37,39^ Although this approach has been proposed for over 20 years, it has not been widely adopted for AAV vector production.^40^ One possible reason for its limited use is the need for a large plasmid (~22 kb), posing challenges during manufacturing, particularly in terms of scalability, purification, and stability. Additionally, large plasmids exhibit lower transfection rates and high toxicities compared to smaller plasmids.^41–43^ Recently, another dual-plasmid system, pOXB, which combines the AAV vector genome and *rep*-*cap* sequences together on a single plasmid, and provides AdV helper genes on a separate plasmid.^44^ A third possible dual-plasmid configuration (pLV) is to combine the AAV genome with the AdV helper sequences, while expressing *rep* and *cap* on a separate plasmid. Interestingly, the pOXB design outperforms the pDG-like and pLV dual-plasmid systems in terms of productivity.^44^ Here, we have developed an all-in-one single-plasmid AAV production system called AAVone. In the AAVone system, the AdV helper genes (*E2A*, *E4orf6* and *VA RNA*), AAV helper genes (*rep* and *cap*), and the AAV vector genome are assembled into one compact pAAVone plasmid with a 13-kb backbone. High-quality AAV vectors can be consistently, efficiently, and easily produced by transfecting packaging cells with relatively low amounts of a single plasmid. This makes the AAVone platform a straightforward, cost-effective, and highly reliable system for AAV production, particularly well-suited for generating GMP-grade AAV vectors.

## RESULTS

### Creation of AAVone single-plasmid system for AAV vector production

To create an efficient single-plasmid system for AAV vector production, it is critical to control the total plasmid size, as it can modulate gene transfer efficiency.^41–43^ Among the three plasmids for AAV vector production, the AdV helper plasmid is the largest. Researchers have continually made efforts to reduce the size of the AdV helper, such as pBHG10, pXX5, pXX6, and pAdDeltaF6.^12,45^ The AdV helper plasmid used in this study, pHelper, is 11.6 kb in size and contains 9.3 kb of the AdV2 genome. We minimized the pHelper plasmid by removing the introns present in the *E2A* and *E4orf6* genes **(Figure S1)**. This modification led to a mini-pHelper that retains only 6.1 kb of the AdV2 genome with a total plasmid size of 8.4 kb **(Figure 1A)**. When co-transfected with the *trans* and *cis* plasmids, the mini-pHelper based system (mTri-plasmid) was capable of supporting AAV vector production, with higher productivity compared with parental pHelper (pTri-plasmid) **(Figure 1B)**.^46^ We then created two versions of dual-plasmid systems that are based on the mini-pHelper: a pDG-like version and a AAVdual (pLV-like) version.^44^ In the pDG-like system, *rep-cap* was assembled into mini-pHelper, resulting in a 13.4-kb plasmid, which is much smaller than the pDG plasmid.^39,40^ In the AAVdual version, the GOI cassette was integrated into the mini-pHelper, resulting in a construct with a 8.4-kb backbone (pAAVdual) **(Figure 1A)**. The AAVdual system exhibited AAV vector yields that were comparable to, or surpassed those of the pTri-plasmid system, whereas the pDG-like system displayed comparable or reduced packaging efficiencies **(Figure 1B)**.

**Figure 1.**
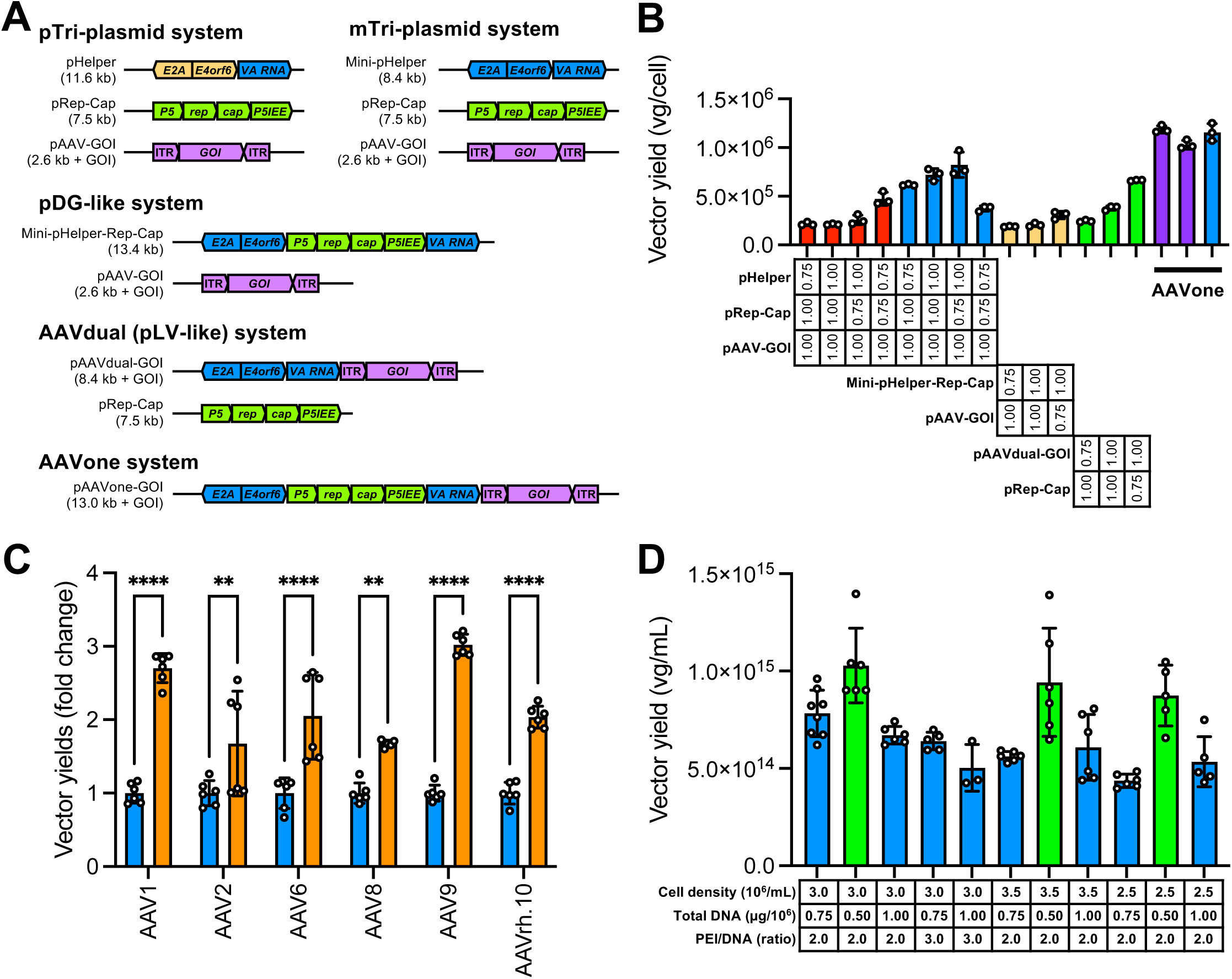
Design and testing of the AAVone production system. **(A)** Schematic illustrations of the AAVone, triple-plasmid (standard, pTri-plasmid; and mini-pHelper, mTri-plasmid) systems, dual-plasmid (pDG-like and AAVdual) systems, and the AAVone system. **(B)** A comparative analysis of AAV productivity using AAVone and alternative systems in adherent HEK-293T cells. For every system compared, the total pDNA amount was maintained at 0.5 µg/well (24-well plate), and the PEImax to DNA ratio was consistently kept at 3:1. The plasmid molecular ratios for each system is displayed under each data set. **(C)** A comparison of vector yields with different serotypes between the AAVone and the pTri-plasmid system in suspension HEK-293T cells. The pTri-plasmid set was transfected at equal plasmid molecular ratios at a total DNA amount of 1 μg per million cells, at a cell density of 3E6 cells/mL. Crude lysate samples were quantified using qPCR with primers targeting Egfp, and data were normalized to the average values of each pTri-plasmid group. **(D)** Productivity optimization of the AAVone system in suspension cultured HEK-293T cells as assessed by crude lysate titration. Different cell densities, total pDNA, and transfection reagent (PEImax) were examined.

Encouraged by the positive outcomes from the dual-plasmid systems, we consolidated all of the *cis* and *trans* essential elements for AAV production into a single-plasmid, pAAVone, with a 13-kb compact backbone **(Figure 1A)**. In this system, the *rep-cap* expression cassette was positioned between *E4orf6* and *VA RNA*, while the GOI cassette was situated between the *VA RNA* and the bacterial origin of replication (*ori*). In this configuration, the *VA RNA* is closer to the left (L)-ITR and bacterial elements are located after the right (R)-ITR. The compact pAAVone plasmids are easy to clone and amplify without reducing yields (**Figure S2**).

With the AAVone system, AAV vectors can be produced by transfecting a single-plasmid into producer cells. For example, AAV2-*CMV*-*Egfp* vectors were generated by transfecting pAAVone-AAV2-*CMV*-*Egfp* plasmid into adherent HEK-293T cells. As shown in **Figure 1B**, the output from the tri- and dual-plasmid systems varied greatly with changes in the proportions of the three plasmids, while the AAVone system produces consistent and higher yields in multiple tests.

When the plasmids were tested by transient transfection **(Figure S3)**, the AAVone system demonstrated comparable levels of EGFP expression (**Figure S3A,B**) and equivalent percentage of positively transfected cells (**Figure S3C**) to those achieved by the pTri-plasmid system. It is important to highlight that in these experiments, the pAAVone-AAV2-*CMV*-*Egfp* construct (15.8 kb) used in the AAVone system and pAAV-*CMV*-*Egfp* construct (5.4 kb) used in pTri-plasmid system carried the same AAV-*CMV*-*Egfp* genome design. In addition, the AAVone and the pTri-plasmid system (with molecular ratio of pHelper:pRep-Cap:pAAV-*CMV*-*Egfp=*1:1:1) used the same amount of total pDNA masses. Under this experimental parameter, the molecular ratio of pAAVone-AAV2-*CMV*-*Egfp* to pAAV-*CMV*-*Egfp* plasmid is 1:0.64. When pAAVone-AAV2-*CMV*-*Egfp* and pAAV-*CMV*-*Egfp* were transfected at the same mass (molecular ratio is 1:2.93), EGFP from the pAAV-*CMV*-*Egfp* plasmid was much stronger than what was produced by transfection with pAAVone (**Figure S3**). This data suggests that the increased AAV productivity of the AAVone system is not due to increased transfection efficiency.

### The AAVone system achieves high AAV vector yields in suspension-cultured cells

To assess the compatibility of the AAVone platform for large-scale manufacturing, we initially evaluated its packaging yields in suspension-cultured HEK-293T cells. AAVone consistently resulted in a two- to four-fold increase in packaging yields compared to the pTri-plasmid system across various serotypes **(Figure 1C and Table 1)**. We next tested various parameters, including different cell densities (ranging from 2.5E6 to 3.5E6 cells/mL), PEI/DNA ratios (ranging from 2:1 to 3:1), and pDNA amounts (ranging from 0.5 to 1.0 µg per 1E6 cells). The AAVone system demonstrated optimal performance when 0.5 µg of pDNA was used for every 1E6 cells, with a PEI/DNA ratio of 2:1, and transfecting at a cell density of 3E6 cells/mL **(Figure 1D)**. The most important parameter for vector production was the pDNA amount. The highest yields were obtained for transfections using 0.5 µg pDNA per 1E6 cells condition.

For high-yield AAV serotypes such as AAV1, AAV5, AAV8, AAV9, and AAVrh.10, the AAVone system easily achieved crude lysate titers exceeding 1E15 vg/L in different production scales without any optimization **(Table S1)**. Among these serotypes, AAV9 demonstrated the highest yields, with crude titers ranging from 1.96E15 to 3.7E15 vg/L and purified titers ranging from 4.22E14 to 1.03E15 vg/L. AAV9 also exhibited the highest fold changes with AAVone, achieving up to a 4-fold increase in yield compared to the pTri-plasmid system. Even with low-yield AAV serotypes, like AAV2, the AAVone system reliably produced between 4.9E14 and 8.9E14 vg/L of crude lysate titers across various production scales, showcasing its consistent performance. Remarkably, the AAVone system demonstrated the ability to generate AAV vectors with transgene genome sizes ranging from 2.2 to 4.7 kb, while the sizes of the corresponding plasmids varied between 15.2 and 17.7 kb **(Table S1)**. Despite a slight decrease in AAV vector yields with increasing transgene sizes, the AAVone system attained 1.94E15 vg/L of crude lysate titer for a 4.7-kb vector packaged with AAV9.

We further evaluated the productivity of the AAVone system in suspension-cultured HEK-293 cells, which are commonly used in GMP-level AAV production. In general, the yields achieved by the AAVone system in the suspension HEK-293 cells were about half of those obtained with HEK-293T cells (**Table S1**). The AAVone system showed adaptability with various transfection agents, yielding high outputs with both PEImax and FectoVIR. Specifically, for AAV9 packaged with a 2.2-kb cassette, AAVone reached up to 1.53E15 vg/L of crude lysate titers with PEImax. With AAV2, the system produced over 4E14 vg/L with PEImax, and more than 6E14 vg/L with FectoVIR. These results collectively indicate that the AAVone system is highly effective across various transgenes, AAV serotypes and cell types, and shows compatibility with different transfection reagents. Moreover, the AAVone system can be easily scaled for production using bioreactors **(data not shown)**.

### AAV vectors generated by AAVone are similar to those produced by tri-plasmid systems

Assessing the quality of AAV vectors is of paramount importance when appraising a novel production system. To directly compare the performance of the AAVone system with tri-plasmid systems, AAV2-*CMV*-*Egfp* vectors were produced across all platforms using suspension-cultured HEK-293T cells. A total of 0.75 µg/1E6 cells of pDNA was used for this examination. Vectors were isolated at specific refractive index (RI) values, ranging from 1.369 to 1.371, following two rounds of CsCl gradient ultracentrifugation. The composition of the capsid proteins, VP1, VP2, and VP3 showed an approximate 1:1:10 ratio across the three systems, with no observable differences between the vectors **(Figure 2A)**. Purified vector DNA also showed that the predominant species were the expected 2.7-kb full-length genomes (**Figure 2B**). All three preparations indicated the presence of truncated or partial genomes with no visually discernable differences in the sizes and abundances of the partial genomes. It is notable that the Tapestation software identified additional low-molecular weight species under 500 bp in size for the pTri- and mTri-plasmid systems, but these species were not identified in the AAVone system (**Figure 2B**). Mass photometry analyses also did not reveal major differences in particle masses between vectors produced by the three platforms (**Figure 2C**). We next evaluated whether there were differences in the transduction profiles between vectors produced by the three systems. Transduction of HeLa cells indicated no clear differences in EGFP expression at 48-hrs post-transduction (**Figure 2D**).

**Figure 2.**
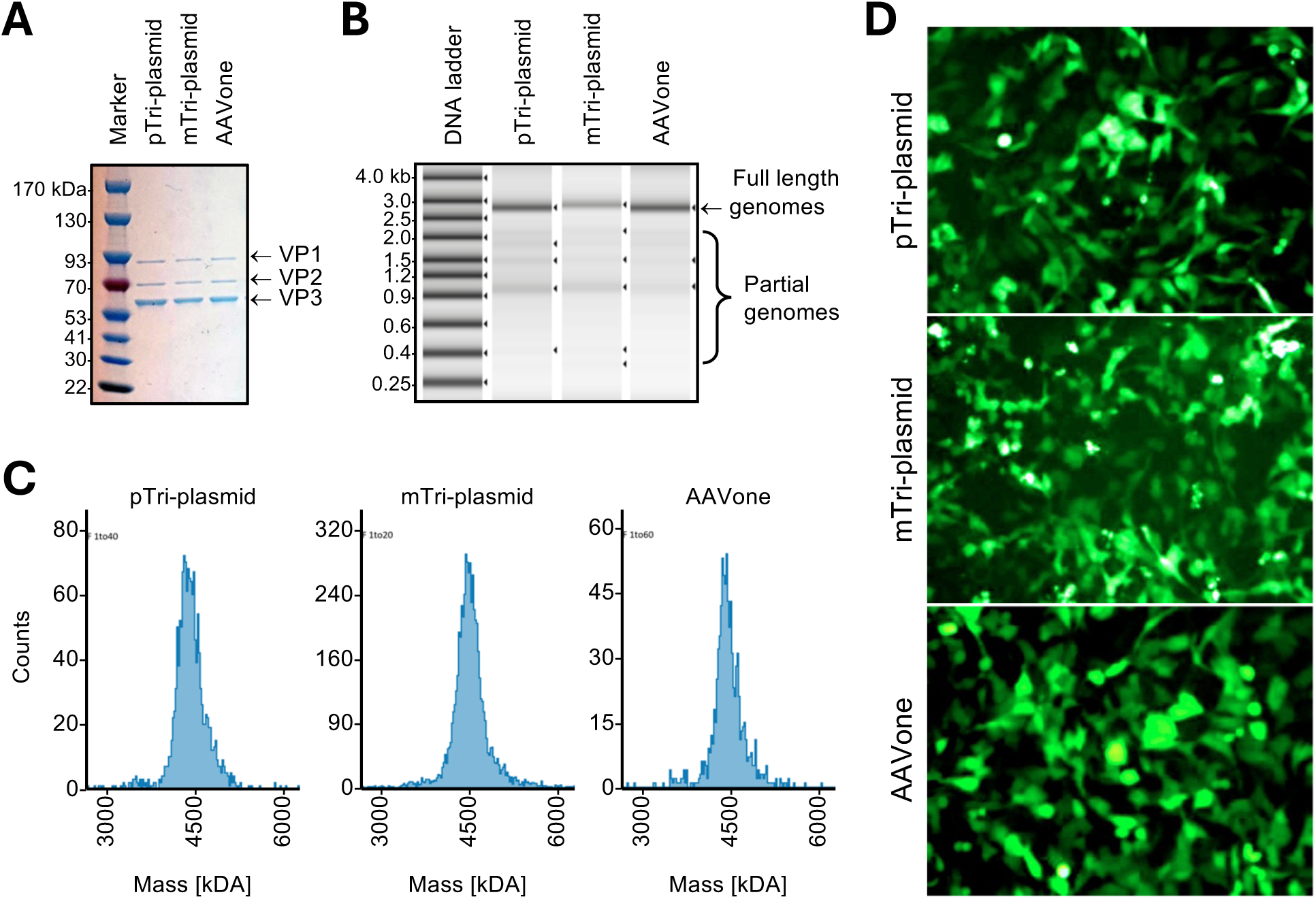
Characterization of AAV vectors produced by the AAVone and tri-plasmid systems. Purified AAV vectors produced in suspension HEK-293T under different production platforms. **(A)** Purified AAV vectors (~5E11 particles) from pTri-plasmid, mTri-plasmid, and AAVone were electrophoresed and visualized by Coomassie G-250 staining. **(B)** Automated electrophoresis analysis (TapeStation) of vector DNAs illustrating full-length genomes (arrow) and partial genomes (bracket). Arrow heads designate specific bands identified by software. **(C)** Mass photometry analysis of AAV particles. y-axis, particle counts; x-axis, mass of particles. **(D)** Transduction activity of vectors packaging the AAV2-*CMV-Egfp* reporter produced by AAVone or triple-plasmid systems in HeLa cells at an MOI of 10,000 vg/cell. and EGFP expression was observed 48 hrs post-transduction.

### Vectors packaged by the AAVone system and tri-plasmid systems exhibited similar retinal transduction profiles

Retinal tissues are among the most prominent targets for AAV vector-based gene therapies being tested in preclinical and clinical trials; especially for platforms that are based on AAV2. Vectors packaged with AAV2-*CMV-Egfp* using the AAVone, mTri-, and pTri-plasmid systems were administered to mouse retinas by intravitreal (IVT) injection at 1E9 vg/eye. At four-weeks post-injection, we conducted funduscopy imaging to assess EGFP expression and distribution across the retina. We found that administration of the three vectors led to similar levels of EGFP expression and distribution (**Figure 3A**). At six-weeks post-injection, retinas were collected, cryo-sectioned, and stained with anti-EGFP antibody (green), peanut agglutin (PNA, cone photoreceptor marker, white), and DAPI (blue). We observed comparable levels of EGFP and PNA signal overlap across all three vector groups (**Figure 3B**) and similar percentages of EGFP-positive photoreceptor outer segments per section (56-59%) (**Figure 3C**). This finding suggested that vectors produced by the AAVone system performed similarly to those produced by the tri-plasmid systems. To quantify the transduction of the vectors produced by the three systems, we extracted bulk DNA and RNA from the harvested retinas to measure the presence of vector genomes and transgene transcripts. Using digital droplet PCR (ddPCR) we found that the vector packaging systems did not influence transduction (**Figure 3D,E**).

**Figure 3.**
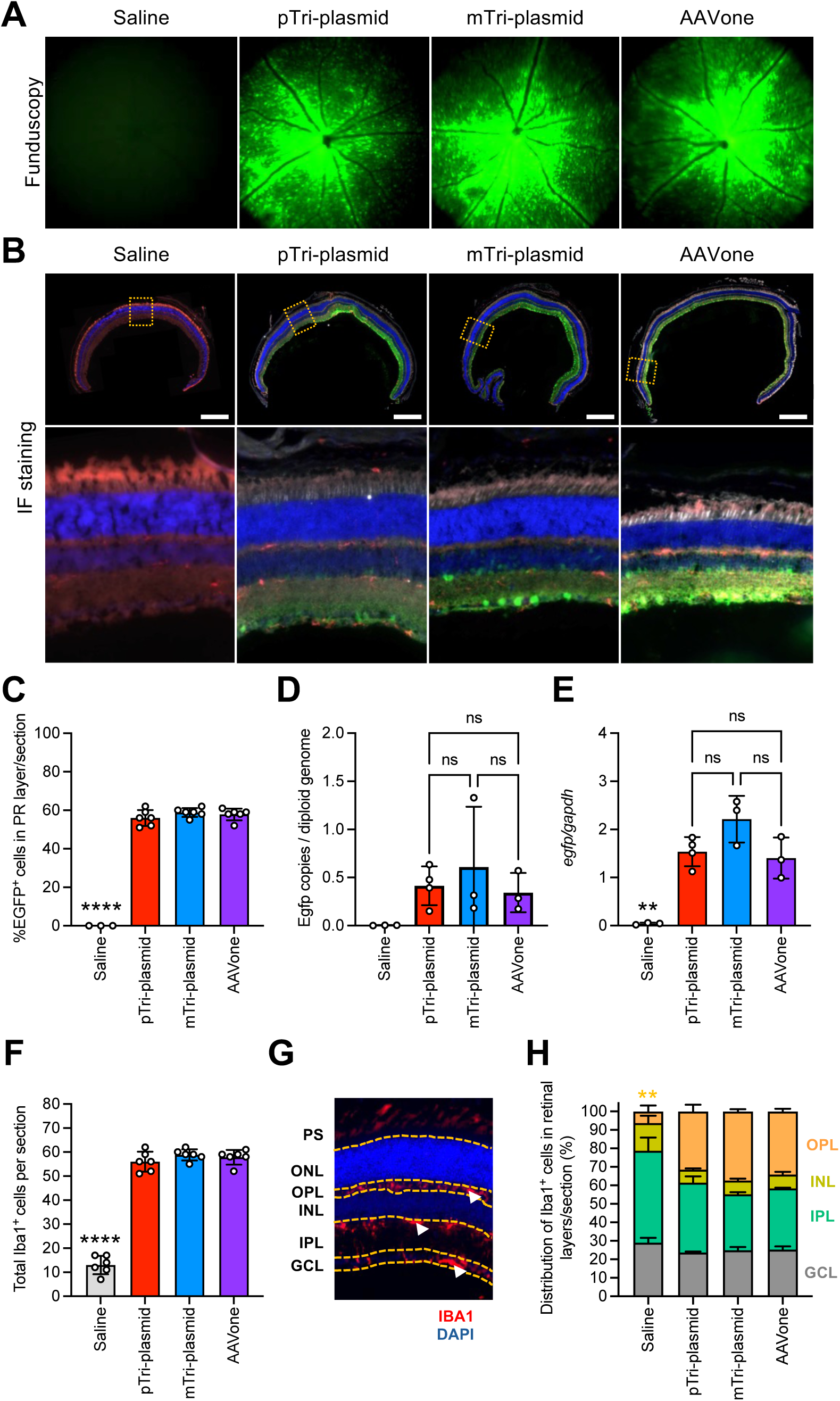
AAVone-packaged AAV2 vectors exhibit similar in vivo efficacies as vectors produce d by triple-plasmid systems. **(A)** Representative funduscopy images of murine retinas at 4 weeks post-injection. **(B)** Representative retina cross-sections stained to visualize EGFP (anti-EGFP, green), nuclei (DAPI, blue), cone photoreceptors (PNA, white), and microglia (IBA1, red) at 6-weeks post-injection. Cross sections of representative whole eye cups (top row) and zoom ins to show transduction of retinal layers (bottom row). Dotted box in top panels show position of zoom-in areas. Scale bars = 0.5 mM. **(C)** Quantification of EGFP^+^ cells in the photoreceptor (PR) layer. **(D,E)** ddPCR quantification of vector genomes **(D)** and vector transcripts **(E)** in retina samples at 6-weeks post-injection (n=4). **(F)** Counts of total microglia in retina cross-sections at 6 weeks post-injection. **(G)** Representative cross-section of a vector-treated retina illustrating microglial (IBA1^+^) infiltration in retinal layers: photoreceptor segment layer (PS); outer nuclear layer (ONL); outer plexiform layer (OPL); inner nuclear layer (INL); inner plexiform layer (IPL); ganglion cell layer (GCL). IBA1^+^ cells (red) and DAPI (blue). **(H)** Percentage of IBA1^+^ cells in each retinal layer. Values represent means ±SD, \*\**p*<0.01; \*\*\*\**p*<0.0001, ns = not significant; by one-way ANOVA and Tukey’s multiple comparison

We next aimed to assess whether vectors produced by the three AAV packaging systems elicit differences in immune responses. It has been well documented that following IVT injection of AAV vectors, microglia are activated and migrate to regions of pathogen-associated molecular pattern (PAMP) activation.^47–50^ We therefore stained cryosections of treated retinas to mark microglia (anti-IBA1). Cells positive for IBA1 were found to increase in the retinas treated by AAV vectors (**Figure 3F**). Notably, we found similar counts of Iba1-positive cells per section across all three treatment groups. We also examined the distribution of Iba1-positive cells infiltrating into the deeper layers of the transduced retinas (**Figure 3G,H**). All three vectors exhibited equivalent stimulation of IBA1-positive cell infiltrates across the retinal across the outer plexiform layer (OPL), the inner nuclear layer (INL), the inner plexiform layer (IPL), and the ganglion cell layer (GCL).

### Vectors packaged AAV1, AAV8, AAVrh.10, and AAV-PHP.eB using the AAVone system show similar *in vivo* transduction profiles as vectors produced with tri-plasmid systems

We next aimed to determine whether the AAVone system was also compatible with other serotypes. We packaged the same ssAAV-*CMV-Egfp* transgene as above with AAV1, AAV8, AAVrh.10, and AAV-PHP.eB capsids by the AAVone, pTri-, and mTri-plasmid systems. Since AAV1 and AAV8 are known to be tropic to muscle, we injected the AAV1- and AAV8-packaged vectors into the tibialis anterior (TA) muscle of adult female mice via intramuscular (IM) injection. Animals were sacrificed at week 2, for those injected with AAV1 vectors; and at week 8, for those injected with AAV8 vectors. TA muscles were harvested and cryosectioned to assess differences in transduction efficiency between the three vectors systems. Fluorescence microscopy of TA cross-sections revealed similar intensities and distributions of EGFP throughout the TA across the three treatment groups for AAV1 and AAV8 (**Figure S4A**) vectors. Additionally, we assessed the immunogenic profiles of vectors produced by the AAVone system compared to vectors produced by pTri- or mTri-plasmid systems. Two-weeks post-injection, there were no observable differences in cellular infiltrates of the TA muscle across the treatment groups (**Figure S4B**).

Both AAV-PHP.eB and AAVrh.10 are well known for their neurotropic properties. We therefore injected adult male C57BL/6J mice at a dose of 1E14 vg/kg by intravenous administration. Two-weeks post-injection, animals were euthanized and brains were harvested. Immunohistochemical staining of brains cryosections showed that AAV-PHP.eB and AAVrh.10 exhibited different patterns of EGFP-positive cells, as expected (**Figure S5**); AAV-PHP.eB primarily transduced neurons, while AAVrh.10 transduced both neurons and glial cells. Importantly, there were no significant differences in EGFP transduction between vectors produced by the three packaging systems among the AAVrh.10 and AAV-PHP.eB groups.

### The AAVone system produces vectors with slightly reduced genome heterogeneity

We next assessed the heterogeneity of the vector genomes using single molecule, real-time (SMRT) sequencing and AAV genome population (AAV-GPseq) analyses.^11,28,51^ Using this approach, we were able to analyze individual AAV vector genomes at a single molecule level. The majority of the reads obtained by SMRT sequencing spanned the entirety of the ssAAV-*CMV-Egfp* reference genome (**Figure S6**). Truncated genomes with snap-back configurations^52,53^ were also observed as reads that only partially spanned the reference from ITR to ITR. These species appear to be similar across the three production platforms, suggesting that there were no substantial differences in truncation events between the vectors analyzed. Instead, the major truncations appear to be centered at the woodchuck hepatitis virus post-transcriptional regulatory element (*WPRE*) (**Figure S5**). We next aimed to assess whether the AAVone system produced vectors with different amounts of non-unit length genomes. The lengths of the reads mapping to the references were tabulated and graphed (**Figure 4A**). The vector genomes from the two tri-plasmid systems had nearly identical read-length distributions, with a major peak at 2,770 nt and multiple smaller peaks at 347 nt, 591 nt, and 1,878 nt. The 2,770-nt population represents the intact AAV genome size, plus two intact 145 nt ITRs, suggesting at ITR repair.^54^ Interestingly, the vector produced by the AAVone system displayed a predominant read length peak at 2,770 nt, with some minor peaks at other lengths. The AAVone system exhibited a slightly higher percentage of intact genomes (58.77%), compared to the pTri-plasmid (51.34%) and the mTri-plasmid (51.41%) systems (**Figure 4A**). Analyses of read alignment start and end positions also indicate a clear reduction in truncation hotspots with the AAVone system when compared to the two tri-plasmid systems (**Figure 4B**). As noted before, the major truncation hotspot appears to be centered at the *WPRE*. Additional truncations are revealed to be centered at the *hGH* polyA sequences (**Figure 4B**).

**Figure 4.**
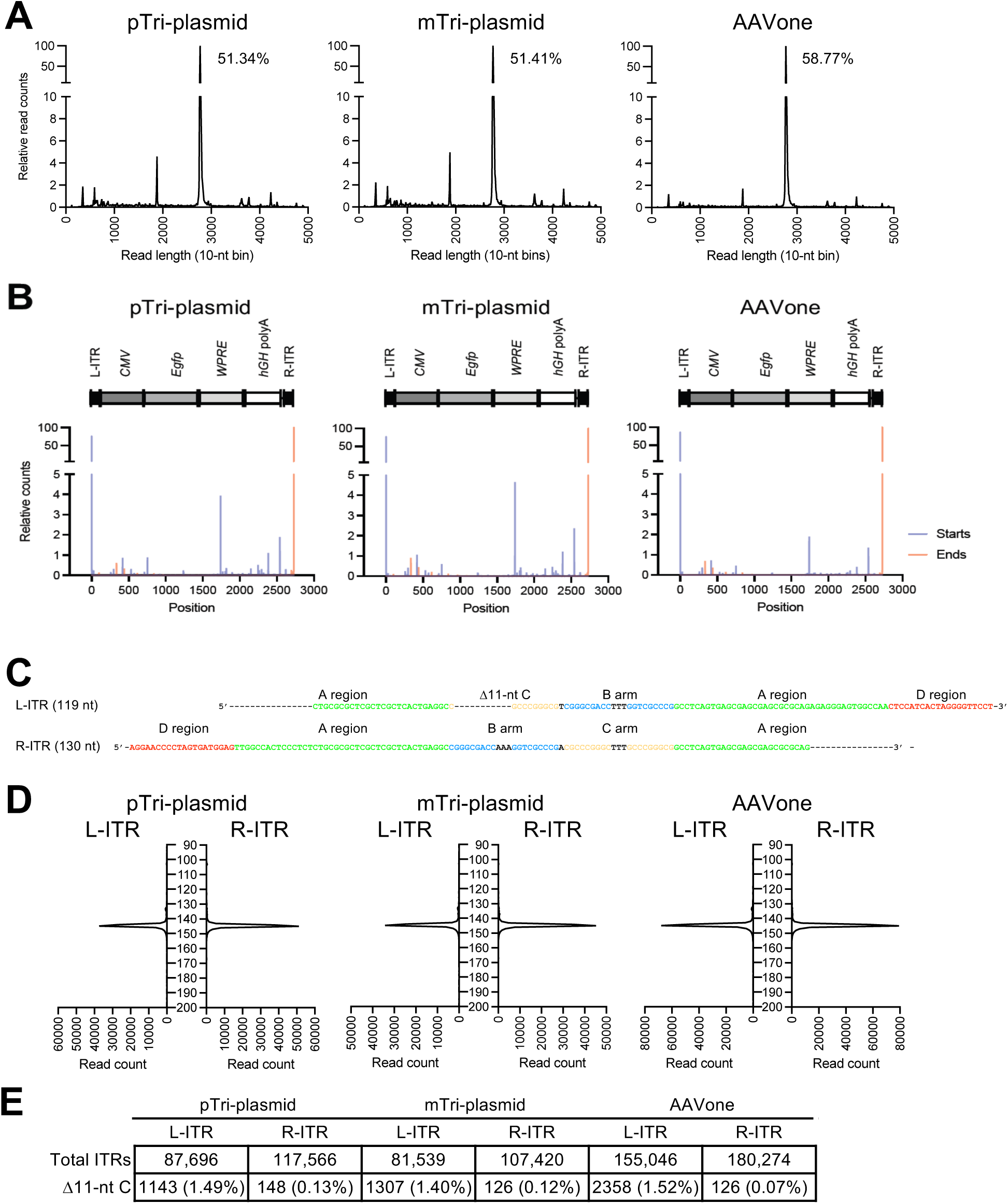
Characterization of AAV vectors produced by the AAVone and tri-plasmid systems. **(A)** Distribution of SMRT sequencing read lengths from vectors produced by pTri-plasmid system (left), mTri-plasmid(center), and AAVone (right). Read lengths are binned into 10-nt increments. The percentage of full-lengh genomes (2,770 ±10 nt) are shown for each graph. The y axes shown are separated into two segments to highlight the low prevalence of truncated genomes. **(B)** Relative counts of mapped read start (blue) and end (red) positions, indicating truncation hotspots. The y axes shown are separated into two segments to highlight the low prevalence of start and end positions beyond the ITR ends. **(C)** Sequences of the 119-nt left ITR (L-ITR) and the 130-nt right ITR (R-ITR) used in the plasmid vector designs. **(D)** Length distribution of the L-ITRs and R-ITRs from the three production systems examined. Read counts are on the x axes, and the read lengths are on the y axes. **(E)** A summary table of the number of 11-nt C arm deleted ITRs at the L-ITRs and R-ITRs identified among the SMRT reads. The percentages of the Δ11-nt C arm ITRs are also shown.

Intact ITRs in AAV vectors are fundamental for transgene expression, and for the overall success of the gene therapy. wtAAV2 ITRs are 145-nt in length and can take on two configurations, flip and flop ^55^. Packaged vectors that undergo proper rolling-hairpin replication show equal distribution of flip and flip at both ITRs. The AAV GOI cassette used by the different production systems are the same. Within the plasmid construct, the R-ITR is 130 nt long, which is 15 nt shorter than the full-length 145-nt ITR. The L-ITR is 119 nt in length and differs from the L-ITR by an 11-nt deletion in the C arm **(Figure 4C)**. Both ITRs were designed in the flop orientation. We found that the ITRs in all AAV vectors were 145 bp in length at both L- and R-ITRs, **(Figure 4D)**. The majority of reads obtained from vectors produced by the two systems were fully aligned to the reference ITRs. Flip and flop are the two major configurations observed in both L- and R-ITRs in AAVone and tri-plasmid systems. They occupy over 90% of the population and display similar flip:flop ratios **(Figure S7)**. Other mutated and truncated forms were also observed in both L- and R-ITRs but they were in the minority. The presence of ITRs missing the 11 nt in the C arm constituted only ~1.5% of the total genomes, implying that the deletion was largely repaired during replication (**Figure 4E**).^56^ Taken together, AAV vectors produced by the AAVone system have nearly identical genomic compositions as compared to those produced by the tri-plasmid systems.

### The AAVone system exhibits alternative plasmid-related impurities in the AAV vectors

Residual input pDNAs are one major concern for AAV production. In the tri-plasmid systems, contaminants mainly originate from the backbone of the *cis* plasmid, while *trans* plasmid sequences are packaged at lower levels.^29^ One major difference between the AAVone and the tri-plasmid systems is the oversized 13-kb backbone, which not only includes the bacterial *ori* and the antibiotic selection gene (*kanR*), but also contains *rep*, *cap*, *E2A*, *E4orf6* and *VA RNA*. In order to examine the impurities originating from plasmids found in vectors produced by AAVone and tri-plasmid systems, we aligned SMRT reads to the AAV-GOI plasmid references. We found that the two ITRs were where the majority of the reads mapped **(Figure 5A-C)**. We also showed that the reads mapping to the plasmid backbone tend aggregate proximal to the ITRs, with decreasing coverage when sequences are further away from the ITRs. Within the AAVone system, *VA RNA* constituted 2.35% of the total sequences **(Table S2)**, making it the most prevalent contaminant. The next most abundant type of contaminant was the plasmid bacterial backbone (0.87%). Other elements, such as *E2*, *E4*, *rep,* and *cap*, which are more distant from the ITRs, exhibit considerably lower percentage of mapped reads **(Table S2)**. The dominant contaminant in vectors produced by the tri-plasmid systems originated from the *cis* plasmid backbone (3.95% in the pTri-plasmid and 2.69% in the mTri-plasmid system). These observations were validated through qPCR analysis focusing on various regions of the pAAVone plasmid **(Figure 5D)**. Thus, the dominant contaminants in AAVone-produced vectors were different from the contaminants related to vectors produced by the tri-plasmid systems.

**Figure 5.**
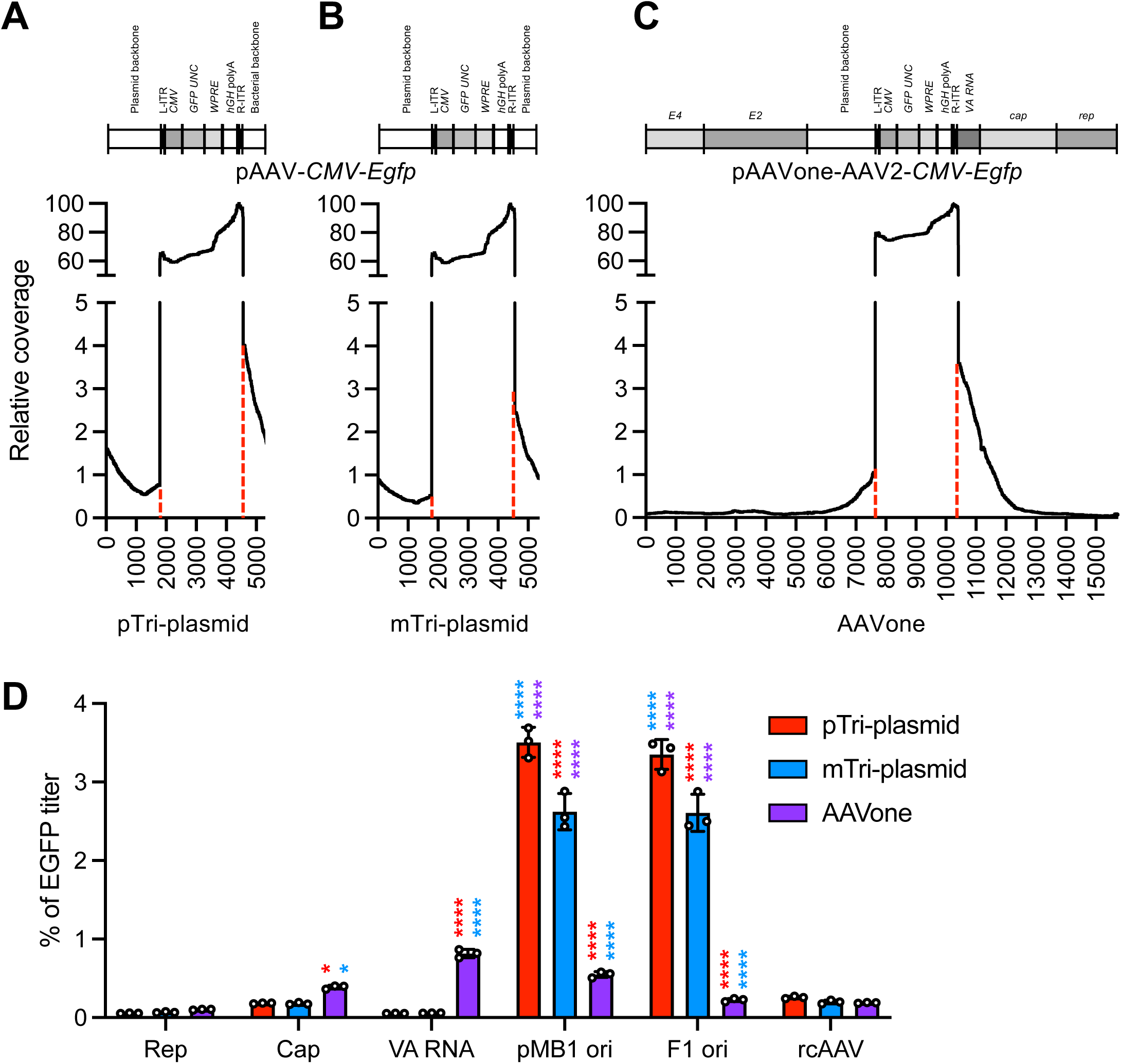
Abundance of producer plasmid contaminants in AAV vectors generated by triple-plasmid systems and AAVone. **(A-B)** Coverage traces of aligned SMRT reads to the pAAVone **(A),** pAAV-CMV-EGFP **(B)** and AAVone **(C)** plasmid references. Dashed lines indicate the boundaries of the ITR references. **(D)** ddPCR quantification of *trans* plasmid sequences (*rep* and *cap*), AdV helper plasmid (*VA RNA*), backbone sequences (pMB1 *ori* and *F1 ori*), and rcAAV in vector preparations. Values represent means ±SD (n=3). *, *p*≤0.05; ****, *p*≤0.0001 by 2-way ANOVA. Color of astrisks correspond to *p* values when compared to pTri-plasmid (red), mTri-plasmid (blue), and AAVone (purple) data.

hcDNA contaminants originating from the packaging cell line used in vector production represent another form of genomic contaminant that may impact the safety of gene therapy vectors. Overall, there was no notable difference in hcDNAs among the AAV vectors produced by the three production systems, with each showing approximately 3% of the total reads **(Table S2**, **Figure S8)**. In all of the reads that mapped to the hg38 genome, the average (~1,300 nt) and mean (~1,140 nt) read lengths were very similar among the three systems **(Figure S9)**. These read lengths are shorter than the smallest oncogene size (1,664 bp).^57^ The number of unique reads spread across each chromosome, suggested at an overall random distribution among all three AAV samples **(Figure S8).**^27,28^ Interestingly, reads mapping to mitochondrial DNA (chrM) appeared to be enriched at the D-loop^27^ **(Figure S10).** However, chrM contaminants that were chimeric with the GOI were also rare (data not shown). Among all of the hcDNA-mapped reads, irrespective of the production system, only 15% appear to be chimeric genomes, hcDNAs recombined with vector genomic sequences **(Table S2**, **Figure S8).** However, when these chimeric reads were mapped back to the GOI reference, and visualized, the majority of reads mapped imperfectly with the hGH polyA **(Figure S11)**. These results indicated that the chimeric reads were predominantly false positives.

In summary, our SMRT sequencing results demonstrate that the AAVone system exhibits significantly lower levels of bacterial backbone sequences, comparable or slightly higher *rep, cap*, *E2*, *E4, and hcDNA* contaminations, and a increase in *VA RNA* sequences compared with the tri-plasmid systems. Despite this shift in the nature of contaminants, the overall total number of genomic contaminations for the AAVone system were decreased (**Table S2**).

### The AAVone system does not pose a risk for generating rcAAV contaminants

Another concern for the AAVone system is the potential for generating replication-competent AAV (rcAAV). The FDA requires that AAV products be screened for the presence of possible rcAAVs. Current vector production systems in use have been optimized to reduce rcAAV to levels ≤1 rcAAV per 1E8 vector genomes.^24^ In tri-plasmid systems, rcAAVs arise through the recombination of DNAs carrying ITRs with *rep* and *cap* sequences originating from the pRep-Cap plasmid.^58^ In the AAVone system, *rep*, *cap,* and the ITRs coexist within the same plasmid. A cell-based rcAAV assay performed by an external contract research organization indicated that rcAAV levels were below one per 1E9 vector genomes for AAV vectors generated by the AAVone system. After three rounds of amplification with AdV, all three flasks in the 1E9 vector genome group showed no presence of rcAAV, and only one out of three samples tested positive for rcAAV in 1E10 vector genomes. This underscores the AAVone system can effectively produce safe AAV vectors with low rcAAV. Using qPCR primers targeting both ITR and *rep* to detect ITR-*rep* hybrids.^59^ which represent a potential for rcAAV contamination, comparable levels were observed among the AAVone and tri-plasmid systems (**Figure 5D**). Analyses of the SMRT reads representing genomes obtained from vectors produced by the AAVone system showed that the majority of reads mapping to the *rep* or *cap* genes also co-mapped with the GOI sequence (89 out of 99 (89.9%) and 201 out of 230 (87.39%), respectively). It is noteworthy that in the tri-plasmid systems, reads that map to either *rep* or *cap* from the *trans* plasmid, which inherently lack ITRs, also displayed a high degree of mapping to ITRs **(Table S2)**. For a read to represent an rcAAV genome, it has to contain the ITRs, *rep*, and *cap* in one sequence.^58^ Our analysis revealed that among the vector genomes from the AAVone system, such chimeric reads were present in 90 out of 196,753 reads (0.045%) **(Figure 6)**, which was slightly higher than those of the pTri-plasmid system (27 out of 131,989; 0.021%) and mTri-plasmid system (22 out of 119,029; 0.018%). Nevertheless, among all of these tri-chimeric sequences, none contained an intact ITR-*rep-cap*-ITR configuration **(Figure 6)**.

**Figure 6.**
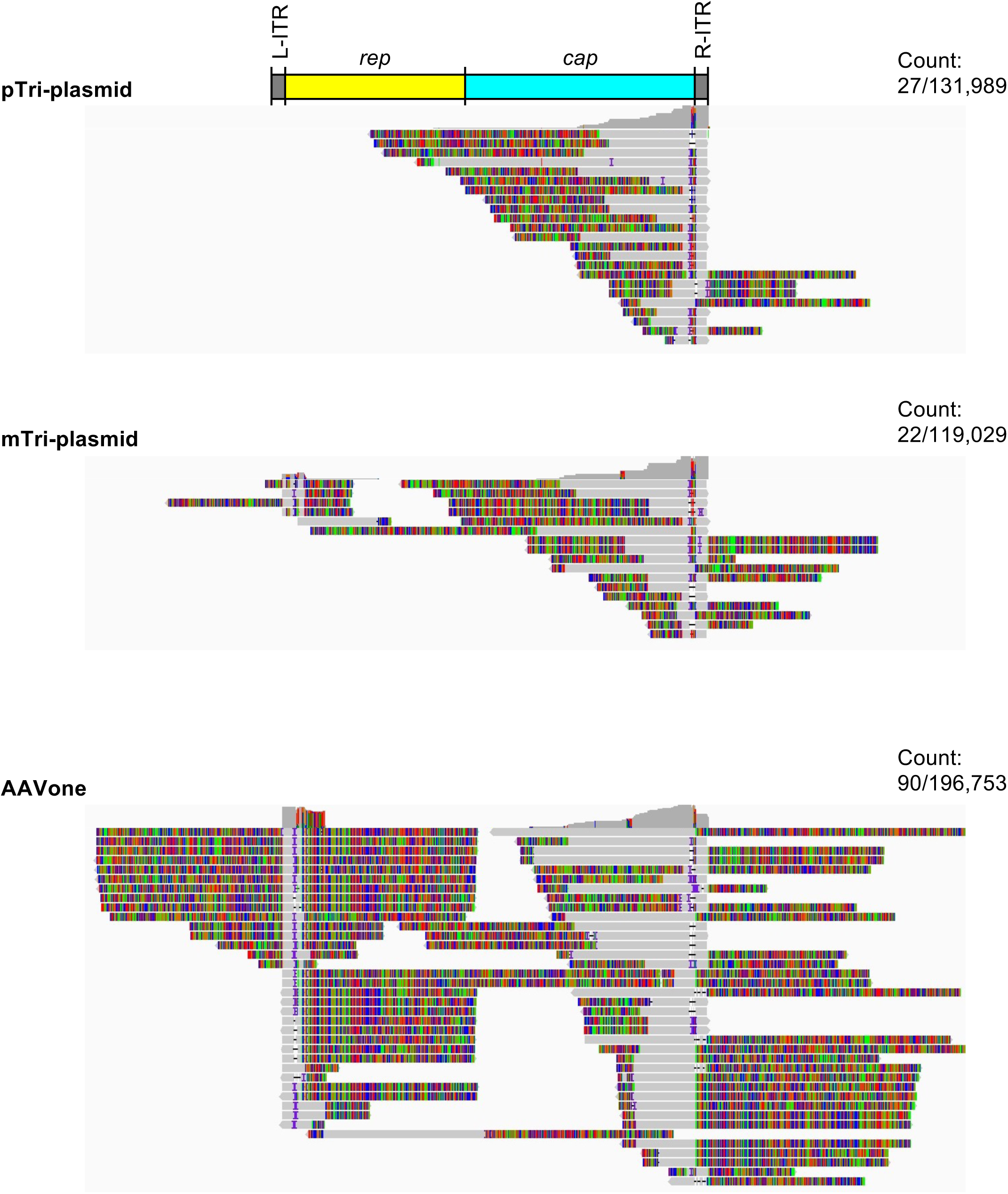
Examination of SMRT reads for the presence of rcAAV. IGV display of SMRT reads representing genomes obtained from vectors produced by pTri-plasmid (top), mTri-plasmid (middle) and AAVone (bottom) systems. Alignments are shown with soft-clipped bases visible. Read matches (gray), mismatches (colored), and insertions/deletions (speckles) are shown.

### The AAVone system enhances production and full-particle ratios while reducing impurities using less plasmid DNA

In the tri-plasmid systems, a relatively high amount of input plasmids (1 µg/1E6 cells) is typically needed to achieve high AAV vector yields.^60–63^ Surprisingly, the AAVone system allows for a substantial reduction of input plasmid, while still increasing AAV yield in both HEK-293T and HEK-293 cells **(Figure 7A and 7B).** In HEK-293T cells, using plasmids of 15.2 kb or 15.8 kb carrying *CMV-Egfp* genomes, the AAVone system reached its highest yield at 0.25 µg/1E6 cells, a result that was independent of the serotype **(Figure S12)**. In HEK-293 cells, the optimal plasmid amounts were between 0.25 to 0.50 µg/1E6 cells **(Figure 7A)**. The reduced input plasmid also increased transfection efficiency for AAV2 vectors (**Figure S12A**). The use of reduced plasmids slightly increased cell viability for both AAV2 and AAV9 vectors **(Figure S12B),** while cell densities reached up to 6E6 cells/mL (**Figure S12C**). Overall, the optimal plasmid dose required for the AAVone system is approximately one half to one fourth of the amount typically utilized in tri-plasmid systems.

**Figure 7.**
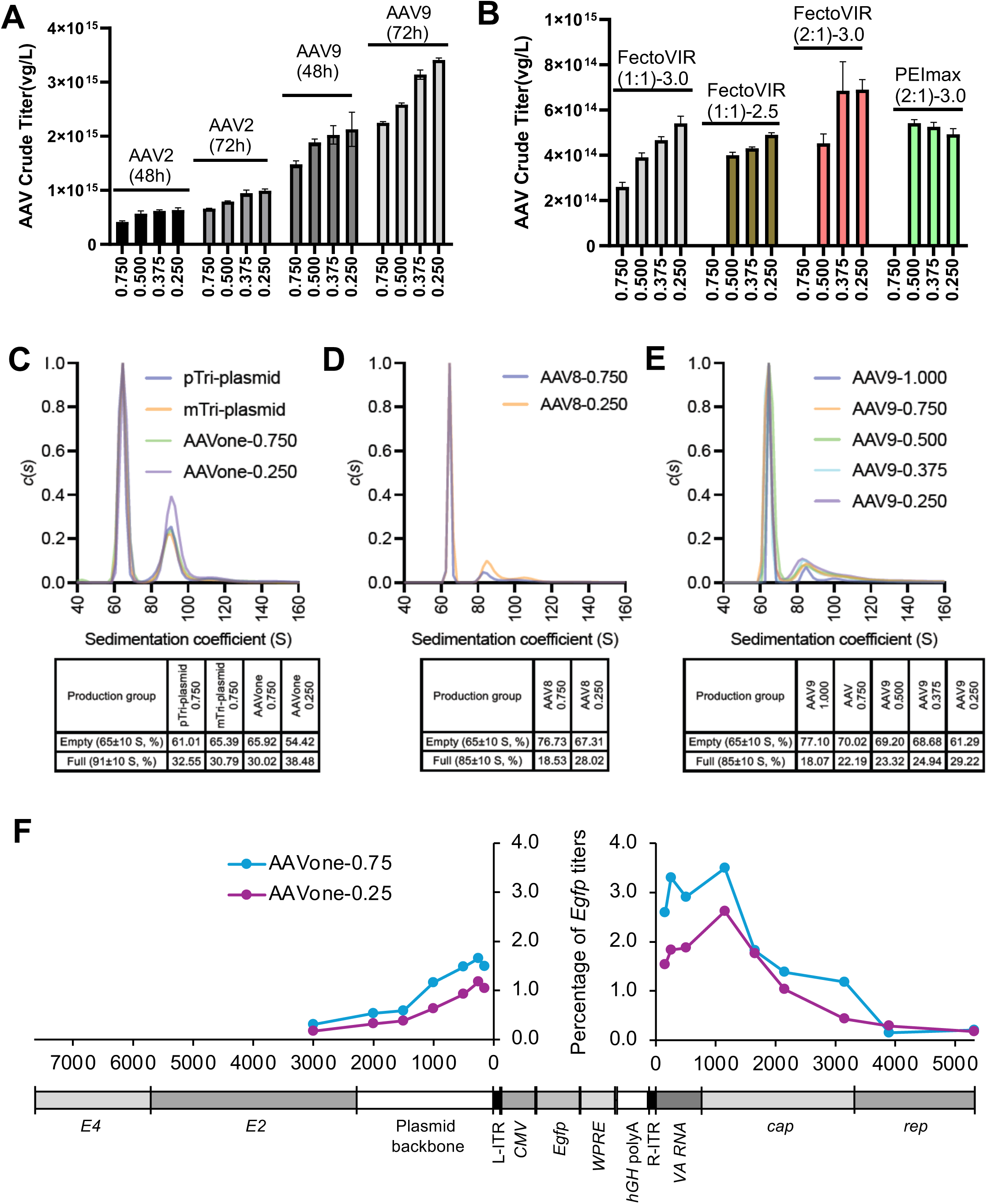
Reduction of AAVone plasmid increases the quality of vectors. **(A)** and **(B)** Effect of pDNA amount on AAV productivity of suspension HEK 293T(A) and HEK 293 (B) cells. **(C-E)** Analytical ultracentrifugation analyses of AAV2 vectors produced by triple-plasmid transfections (pTri-plasmid and mTri-plasmid) and AAVone produced under different plasmid amounts **(C)**. AUC analyses of AAV8 **(D)** and AAV9 **(E)** vectors under different plasmid amounts x axis = sedimentation coefficient (S). The y axis = normalized distribution *c*(*s*). **(F)** qPCR analyses of DNase-resistant target regions packaged with AAVone-produced vectors. x axes represent positions moving away from the ITRs.

The use of reduced input plasmid with the AAVone system has additional benefits. It led to increased full-to-empty capsid ratios in the whole production system (**Figure 7C-E**), which are desirable for obtaining higher-quality AAV vectors with a larger proportion of functional, transgene-carrying capsids. When utilizing the same quantity of input pDNAs to produce AAV2 vectors (0.75 µg/1E6 cells), the AAVone system exhibited a similar full-to-empty ratio as compared to what was achieved by the tri-plasmid systems (**Figure 7C**). All systems demonstrate two prominent peaks at 66 S and 92 S, which correspond to empty AAV2 particles and full capsids with 2.8-kb genomes, respectively. Approximately 30% of AAV particles found in crude lysates for each production system were full capsids. When the plasmid was reduced to 0.25 µg/1E6 cells with the AAVone system, the full particle ratio rose from 30.02% to 38.48%. Similarly, titers for AAV8 and AAV9 exhibited a ~10% increase in full AAV particles when using a reduced amount of input pDNA (**Figure 7D,E**).

The use of reduced pDNA with the AAVone system can also help minimize plasmid-related impurities. Reducing the pDNA amount led to a decrease in plasmid impurities on both sides of the ITRs **(Figure 7F)**. In summary, reducing the input plasmid increases packaging efficiency, leads to increased full-to-empty particle ratios, and reduced plasmid-related impurities.

## DISCUSSION

### Advantages of the AAVone system

The AAVone system described here is designed to bring all the essential components for AAV production into a single-plasmid (**Figure 1**), while retaining all of the essential vector performance features and qualities expected of those produced by standard tri-plasmid transfection platforms (**Figures 2 and 4**). These features include the preservation of important characteristics such as capsid content, genome and ITR integrity, transduction efficiency, ratio of full-to-empty particles. With this system, users can integrate their existing GOIs into the AAVone manufacturing process without the need to develop additional upstream and downstream processes, analytical release, or characterization assays.

A notable benefit of using the AAVone system is the considerable reduction in costs associated with raw materials, as it only uses a small quantity of a single-plasmid, instead of three plasmids. Plasmids represent one of the most expensive components in AAV vector production. The large-scale production of three GMP grade plasmids with over 95% purity and absent of process-related impurities, continues to be a significant challenge. Although numerous researchers have dedicated efforts to developing dual-plasmid transfection systems for AAV vector production,^37,39,40,44^ this study stands out as the first to investigate the production of AAV vectors using a single-plasmid. The system enables the production of AAV vectors using only about one-quarter to one-half of the total pDNA normally required with conventional approaches **(Figure 7)**. Lowering the input of pDNA not only reduces costs, but also leads to decreased genomic contaminants, increased full-to-empty capsid ratios, and increased titers (**Figure 7**). Combined with a two- to four-fold increase in AAV vector productivity, the AAVone system uses only one-tenth of the total pDNA to achieve the same titers produced from the pTri-plasmid system. The AAVone system’s single-plasmid approach also addresses the challenges faced with traditional tri-plasmid systems, such as obtaining a consistent vector product **(Figure 1B** and **Table S1)**. Finding the optimal ratio of the three plasmids, maintaining plasmid stability, and ensuring consistent copy numbers across different production batches is complex and often times challenging when considering the different cell lines, AAV serotype, and the size and composition of the rAAV cassette used.^60–63^ The single-plasmid system minimizes the potential for introducing plasmid-related variation; thereby, enhancing the consistency of the AAV vectors produced.

The AAVone system potentially offers a higher effective AAV vector production efficiency compared to the pTri-plasmid system. The increased productivity of AAVone system is not due to increased transfection efficiency, but increased production efficiency **(Figure S3)**. Theoretically, within the AAVone system, there is a linear relationship between AAV vector production efficiency and transfection efficiency. However, in the tri-plasmid systems, transfection efficiency cannot linearly convert to AAV vector productivity, as only cells that are co-transfected with all three plasmids are able to produce AAV vectors **(Figure S13)**. This effect becomes significantly pronounced at lower transfection efficiencies. It has been reported that at a transfection efficiency of ~60%, only ~7% of cells produce measurable amounts of assembled AAV capsid with the tri-plasmid transfection method, while transfection with the dual-plasmid system increased cells positive for assembled AAV capsid by four- to five-fold.^19^ Moreover, it has been reported that with three different dual-plasmid configurations, pOXB showed significantly superior results compared to the pDG-like and pLV systems.^44^ We also found that the pLV configuration exhibited comparable productivity, and the pDG-like configuration showed a lower production efficiency than achieved with the tri-plasmid method (**Figure 1B**). These findings suggest that co-existence of the AAV vector genome and AAV helper genes may play an important role in the increased vector productivity by AAVone.

### Genomic impurities by the AAVone system

Besides intact GOI genomes, AAV vector production also yields contaminants in the final product, including incomplete AAV vector genomes, and DNAs from the producer plasmids and host genome.^23^ The presence of contaminants in clinical vectors is considered undesirable because it can lead to increased immune responses against the vector, potential integration into the host genome, and other safety concerns.

Different types of packaged incomplete vector genome species are present in AAV vectors,^52^ but most of them are snap-back forms (**Figure S4**), which are double-stranded DNAs containing either 5′ or 3′ ITRs.^23^ The proportion of incomplete genomes are sequence dependent, and regions with high secondary structure such as short hairpin RNA (shRNA) result in a greater proportion of snap-back genomes.^28,52^ In our AAV-*CMV*-*Egfp* constructs, the major truncation hotspot appears to be centered at the *WPRE*, *hGH* polyA, and *CMV* sequences, which are close to ITRs (**Figure 4A,B** and **Figure S6**). In our previous studies, byproduct empty rAAV particles were not truly empty, but contained different types of short vector genomes that were enriched for sequences corresponding to the ITRs.^64^ We were unable to completely remove these aberrant genomes from rAAV preparations, even after extracting thin bands related to full capsid fractions following CsCl ultracentrifugation (**Figure S2** and **Figure 4A,B**). However, the larger size of the pAAVone plasmid may contribute to the reduction of aberrant genomes, as the AAVone system yielded slightly more intact genomes and significantly less the snap-back genomes in the final purified AAV vectors when compared to what is produced by tri-plasmid systems (**Figure 4** and **Table S2**). Furthermore, decreasing the amount of the pAAVone plasmid during production significantly reduced the impurities as well as the empty particles (**Figures 5 and 8**). Both the DNA template switch model and NHEJ mechanism may result in the formation of snap-back genomes.^52^ The use of one large plasmid may overall reduce recombination events between ITR and other plasmid elements, which in turn leads to decreases in genomic impurities. Previous studies have concluded that increasing the size of the plasmid backbone with stuffer DNA can reduce the level of plasmid backbone contaminants, as the two ITRs are not in range of each other to facilitate reverse packaging. The pAAVone plasmid has 13 kb of non-GOI-cassette sequences existing beyond the two ITRs. The oversized design did decrease the abundance of sequencing spanning the bacterial *ori* and the antibiotic resistance gene. All of the high abundance elements were positioned close to ITRs on the plasmid, which supports the idea that the majority of vector product-related impurities are ITR-containing fragments. In addition, the abundance of contaminants is mainly determined by the distance from the ITRs (**Figures 5 and 8D**) and sequences positioned more than 2.5 kb away from the ITRs, such as *rep*, *E2*, and *E4* genes, are rarely packaged into AAV vectors (**Figure 5).** Thus, incorporating genes into one single-plasmid did not significantly increase their related impurities. As the *rep* is located 5.6 kb away from ITR, the pAAVone design did not increase the potential for rcAAV formation. Overall, we found that an oversized backbone design reduced total plasmid-related impurities (**Table S2**).

In conclusion, the AAVone packaging system represents a simple, cost-effective, and highly reproducible AAV production system, particularly suitable for GMP-grade AAV production.

## MATERIALS AND METHODS

### AAV plasmids

All vector constructs were generated using HiFi assembly (NEB), Gibson assembly, or T4 ligation and were scaled up with midi/maxi preparation kits (Takara Bio). The mini-pHelper was generated by deleting introns of *E2* and *E4* genes from the pHelper parent construct. pAAVdual plasmids were cloned by inserting AAV-GOI genomes (including ITRs) into mini-pHelper at the *Pme*I site, located between pMB1 *ori* and *VA RNA*. mini-pHelper-Rep-Cap plasmids were cloned by inserting the *rep-cap* cassette into the *Cla*I site of mini-pHelper, which is located between *E4* and *VA RNA*. pAAVone-AAV2-*CMV*-*Egfp* plasmids were achieved by insertion AAV2-*CMV*-*Egfp* genome into the mini-pHelper-Rep-Cap-AAV2 at the *Pme*I site. pAAVone plasmids with different capsids and transgenes based one pAAVone-AAV2-*CMV*-*Egfp*. All the ITR-bearing plasmids were subjected to restriction enzyme analysis using *Sma*I and/or *Ahe*I, ITR sequencing and/or whole plasmid sequencing to confirm the ITR structures.

### AAV production, purification, and titration

Adherent HEK-293T cells were cultured in DMEM medium with 10% fetal bovine serum (FBS) and 1% penicillin/streptomycin (P/S) in various culture vessels including 24-well plates, 15 cm dishes, or 2 L roller bottles. Suspension HEK-293T and HEK-293 cells were expanded for at least three passages and inoculated into a shake flask with F17 culture medium (Thermo Fisher, Waltham, MA) prior to transfection. The transfections were carried out using either PEImax (Polyscience, Warrington, PA) or FectoVIR (Polyplus, NY), adhering strictly to the provided guidelines. Experimental variations included different cell densities, amounts of pDNA, and ratios of transfection reagent to DNA. Cells were collected for analyses 72 hours after transfection. For initial titer estimations, cell samples underwent a single freeze-thaw cycle and sonication for lysis. Subsequent centrifugation allowed for the collection of supernatants, from which crude titer productivity was assessed. The AAV vectors extracted from these samples were then purified through one or two CsCl-gradient ultracentrifugation steps before undergoing quality evaluation. Prior to titer determination, both crude lysate and purified samples were treated with DNase I to remove un-encapsidated residual DNA, followed by Proteinase K treatment. The quantification of AAV vector productivity was achieved using either Droplet Digital PCR (ddPCR) or qPCR, employing primers and/or probes targeting the vector’s transgene. To establish a standard curve, a plasmid carrying the relevant sequence was linearized. Each experimental batch included a well-characterized AAV vector as an internal standard, ensuring data normalization against this control.

### Mice

Six- to eight-week-old female C57BL/6J mice were administered AAV2-*CMV-Egfp* or saline IVT injection to the left and right eyes at 1E9 GC/dose (n=10 per group). All AAV2 vectors were diluted with saline and fast green to 1.0E9 vg/μL. The IVT injections were performed with glass needles (Clunbury Scientific LLC; Cat no. B100-58-50) to deliver ~1 μL of fluid into the vitreous using the FemtoJet from Eppendorf with a constant pressure of 300 psi and injection time of 1.5 s. At four-weeks post-injection, funduscopy imaging was performed to assess the EGFP expression in the mouse retina. Mice were euthanized at four- and six-weeks post-injection. Eye cups were collected for immune-staining and molecular quantification.

Six- to eight-week-old female C57BL/6 mice were administered AAV1-*CMV-Egfp*, AAV8-*CMV-Egfp*, or saline via intramuscular (IM) injection to the left and right tibialis anterior (TA) muscle. Vectors were diluted with saline to 1E10 vg/µL and administered at 1E11 GC/dose (n=10 per group). At two- and 6-weeks post-injection, TA muscles were harvested and either placed in 4% paraformaldehyde (PFA, for histology) or snap frozen (for RNA and DNA extraction). Following 4% PFA overnight fixation, tissues were sent to UMass Chan Medical School Morphology Core for H&E staining or placed in 30% sucrose for 72 hours before embedding in Tissue-Tek® O.C.T. Compound (Sakura Finetek USA Inc #4583). The embedded tissues were cryosectioned (10 μM slices) and mounted on slides with VECTASHIELD® HardSet™ Antifade Mounting Medium, with DAPI (Vector Laboratories #H-1500-10) for microscopy. All images were visualized with a Leica DM6 Thunder microscope with a 16-bit monochrome camera. Images were processed by LAS X Life Science microscope software.

AAV vectors were intravenously administered to adult male C57BL/6J mice (8–10 weeks old, n=3) at a dose of 1E14 vg/kg per mouse. Two-weeks post-injection, animals were euthanized and intracardially perfused with cold 0.1 M PBS followed by 4% PFA in 0.1 M PBS. The brains were harvested, fixed overnight in 4% PFA, and cryoprotected in 30% sucrose at 4 °C for 2-3 days. Eight series of 40-μm-thick sections were cut using a frozen sliding microtome, collected in sterile Eppendorf tubes containing 30% sucrose in 0.1M PBS, and stored at −20°C until use.

### Immunohistochemistry

Treated retinas were cross-sectioned as previously described ^48^. In brief, eye cups were dissected in cold 1X PBS and fixed in 4% PFA overnight at 4 °C. Cryosections were cut to 12-micron thickness. The following primary antibodies and dilutions were used: chicken anti-EGFP antibody (1:1000; Abcam; Cat no. ab13970), and rabbit anti-IBA1 (1:300; Wako; Cat no. 019-19741). The chromophore conjugate, fluorescein peanut agglutinin (PNA) (1:1,000; Vector Laboratories; Cat no. FL1071) was used to stain cone photoreceptors. All antibodies were diluted in PBS with 0.3% Triton X-100 and 5% bovine serum albumin (CST). Nuclei were counterstained with 4′,6-diamidino-2-phenylindole (1:2000, Sigma-Aldrich; Cat no. 9542). The following secondary antibodies were used: Donkey anti-Chicken IgY (H+L) Highly Cross Adsorbed Secondary Antibody, Alexa Fluor™ 488, A78948 and Goat anti-Rabbit IgG (H+L) Highly Cross-Adsorbed Secondary Antibody, Alexa Fluor™ 594, A-11037. All secondary antibodies were diluted 1:1000 in PBST (PBS with 0.25% tween-20). All images were visualized with a Leica DM6 Thunder microscope with a 16-bit monochrome camera. Images were processed by LAS X Life Science Microscope Software.

Floating brain sections were washed three times with 0.5% Triton X-100/Tris-buffered saline (TBS) and blocked for 30 minutes in a blocking buffer containing 2% BSA in TBS with 0.5% Triton X-100. Primary anti-GFP antibodies were then added, and the sections were incubated overnight at 4 °C. After three washes, the sections were treated with Alexa fluorescent-labeled secondary antibodies in blocking buffer for one hour at room temperature. Following washing and Hoechst counterstaining, brain sections were mounted onto glass microscope slides and cover slipped with Fluoroshield (Sigma Aldrich Cat. #F6182). Tiled confocal images of entire brain sections were acquired and analyzed using the Leica Stellaris-5 confocal microscope system.

### RNA and DNA extractions

Snap-frozen TA muscles were homogenized using Bullet Blender Green Bead Lysis Kit (NextAdvance #GREENE5) in conjunction with a Qiagen TissueLyzer II. Half of the homogenized tissues were processed for RNA extraction using mirVana™ miRNA Isolation Kit (ThermoScientific #AM1561) according to the standard protocol. The remaining homogenized tissues were processed using QIAamp DNA Mini Kit (Qiagen #51306) according to manufacturer’s recommended procedures. Extracted RNAs were reverse transcribed into cDNA using Iscript™ Reverse Transcription Supermix (BioRad #1708841) and Iscript™ gDNA Clear cDNA Synthesis Kit (BioRad #1725035).

Snap-frozen retinas were homogenized using Qiagen 5mm TissueLyser bead (Qiagen #69965) in conjunction with Omni Bead Ruptor Elite. Homogenized retina tissue was processed for RNA and DNA extraction using the Norgen Biotek RNA/DNA/Protein Purification Plus Kit (Norgen Biotek #47700) according to the standard protocol. Extracted RNA was reverse transcribed into cDNA using Iscript™ Reverse Transcription Supermix (BioRad #1708841).

### Digital Droplet PCR

Digital Droplet PCR (ddPCR) was performed on gDNA and cDNA using ddPCR Supermix for Probes (No dUTP), 2X (BioRad #1863024) with the following probe sets: *Egfp*-FAM (Thermofisher #4400291, Assay ID Mr00660654_cn), mouse *Gapdh-*VIC (Thermofisher #4448484, Assay ID Mm99999915_g1), and TaqMan™ Copy Number Reference Assay, mouse, Tfrc-VIC (ThermoFisher #4458366). Input for AL 479, AL 480, and AL 481 ddPCR reactions were as follows: 40 ng gDNA per 20 uL reaction and 0.5 uL cDNA (diluted 1:15) per 20 uL reaction. Droplet generation was performed using a BioRad Automated Droplet Generator using the corresponding manufacturer recommended consumables into a 96-well plate. PCR reactions were conducted on a thermocycler using the following conditions: 95 °C for 10 minutes, 32 cycles of 95 °C for 30 seconds and 61 °C for 1 minute, and a final step of 95 °C for 10 minutes. Droplets were read using a BioRad QX200 Droplet Reader and analyzed using QuantSoft software. Raw reads were exported and *Egfp* values were normalized either to diploid genomes or to housekeeping gene transcripts.

### AUC analysis

AUC was used to quantify empty and full particles. AAV from CsCl index 1.360-1.380 were collected after one round of ultracentrifugation. AUC measurements were performed at 20,000 rpm and 20 °C on a Beckman AUC Optima instrument (Beckman Coulter, Brea, CA, USA) equipped with an An-50 Ti analytical rotor (Beckman Coulter, Brea, CA, USA) using a total number of 150 scans per sample. Analysis of AAV samples by the SEDFIT c(S) model yields a distribution of sedimentation coefficients with each peak in the distribution representing AAV particles with full genomes or empty capsids. Integration of individual peaks yields the sedimentation coefficient (S) and the relative concentration of each species in the distribution.

### Mass photometry

Mass photometry measurements were carried out on a SamuxMP instrument (Refeyn Ltd., Oxford, UK). Prior to each measurement, a calibration was conducted using truly-empty AAV particles, which was produced by transfection of one mini-pHelper-Rep-Cap plasmid in HEK-293T cells.

### qPCR titration plasmid related impurities contaminant amplicons

Purified AAV vectors were treated by DNase I and proteinase K. Vector genome titers and contaminant amplicon titers were assessed by qPCR using SYBR Green. Viral titers were quantified on a StepOnePlus Real-Time PCR System (Applied Biosystems) using the default settings. The procedure involved denaturation at 95 °C for 5 min, followed by 40 cycles of amplification (95 °C for 10 s, 60 °C for 30 s), and concluding with a melting curve. Primers targeting different regions of the pAAVone plasmid are shown in **Table S1**. A plasmid containing the corresponding target sequences was used to make the standard curve. All data were normalized to *Egfp* transgene ct values.

### Characterization of the AAV capsid

The capsid compositions of AAV vectors were determined by Bio-Safe Coomassie G-250 stain. AAV vectors (~5E11 particles) were resolved by electrophoresis on a 10% SDS-PAGE gel and standard Coomassie G-250 stain was performed according to the manufacturer’s procedures (Bio-Rad).

### AAV vector transduction in vitro

The HeLa cell line used in this study were grown in DMEM with 10% FBS and 1% P/S at 37 °C in a humidified environment supplied with 5% CO_2_. For each transduction experiment, ~50,000 viable cells were seeded onto a 24-well plate 24 h before transduction. AAV vectors were added directly to each well at an MOI=10,000. EGFP expression was determined 48 h post-transduction by epifluorescence microscopy.

### Flow cytometry

Cells transfected with the tri-plasmid systems or the AAVone system was collected at approximately 48 h after transfection and processed using a BD Accuri C6 Plus flow cytometer equipped with an autosampler (BD Biosciences, San Jose, CA). At least 50,000 events were collected per sample with the fluidics rate set to slow (14 μL/min) with one agitation/SIP clean cycle between samples. The acquired data were next analyzed with FlowJo software (v10.8; FlowJo, Ashland, OR) using standard gating strategies to select for single cells (based on SSC-A vs. FSC-A and FSC-H vs. FSC-A scatterplots) and to calculate the percentage of GFP-expressing cells (based on FITC-A histogram).

### NGS analysis of AAV vector genomes

NGS of AAV vector genomes was performed by Azenta. The ssAAV viral DNA was extracted using the Invitrogen PureLink Viral RNA/DNA kit following manufacturer’s instructions. DNA samples were quantified using a Qubit 2.0 Fluorometer (Life Technologies, Carlsbad, CA, USA) and sample integrity was checked using Agilent TapeStation 4200 (Agilent Technologies, Palo Alto, CA, USA). Amplicon libraries were prepared using SMRTbell Express Template Prep Kit 3.0. The sequencing library was validated on the Agilent TapeStation and quantified by using Qubit 2.0 Fluorometer. All libraries were combined into a single sample and sequenced across two PacBio Sequel IIe SMRT cells using Sequel II binding kit 3.0 with heteroduplex detection mode on.

### SMRT sequencing and AAV-GPseq

The bioinformatic workflows were conducted using the Galaxy platform.^65,66^ Resulting consensus fastq files were mapped to the references as reported in the study using the Burrows– Wheeler aligner-maximal exact match (BWA-MEM)^67,68^ in PacBio mode (-x pacbio). Aligned reads were visualized with the Integrated Genomics Viewer (IGV) tool version 2.14.0 ^69^ with soft clipping on.

### Statistical Analyses

The results were analyzed by one-way or two-way ANOVA statistical tests were performed in GraphPad Prism 10 along with Tukey’s multiple comparison statistical tests where applicable as described.

## DATA AVAILABILITY STATEMENT

All data are provided in the main text, figures, or the supplemental information, and raw data files are available upon request. Requests for reagents or cell lines used in this study can directed to the corresponding author, Qizhao Wang.

### Declaration of Generative AI and AI-assisted Technologies in the Writing Process

During the preparation of this work the author(s) used ChatGPT to assist with revising the format, grammar, and spelling. After using this tool/service, the author(s) reviewed and edited the content as needed and take(s) full responsibility for the content of the publication.

## ACKNOWLEDGMENTS

G.G. is supported by grants from the UMass Chan Medical School (an internal grant) and by the National Institutes of Health (R01NS076991-01, P01HL131471-05, R01AI121135, UG3HL147367-01, R01HL097088, R01HL152723-02, U19AI149646-01, and UH3HL147367-04). P.W.L.T. is supported by an award from The Bassick Family Foundation, a BRIDGE Fund Award (a UMass Chan Medical School internal grant), and by the National Institutes of Health (1R21AI183080-01A1). W.H. is supported by the National Institutes of Health (R01DA056876)

## AUTHOR CONTRIBUTIONS

R.Y., J.Z., Y.L. designed and performed the AAV productivity experiments; Y.W. and Y.L. conducted all AAV-related characterizations; N.T.T. analyzed the NGS data and prepared figures. M.Cu., T.C., and T.S. designed and carried out animal studies involving AAV1, AAV2, and AAV8, and drafted the parts of the manuscript that encompass this work; X.Y., D.Z., and D.J. conducted animal studies for AAV-PHP.eB and AAVrh.10; Y.L., Z.S., Y.D., L.F., H.H., B.W., and C.X. constructed and prepared all plasmids; L.W., H.Z., Y.L., M.Ch, J.L., and C.C. performed AAV production, purification, and quality checks; G.G. revised and organized the manuscript, and supported the AAV1, AAV2, and AAV8 animal studies; W.H. provided support for the AAV-PHP.eB and AAVrh.10 animal studies. D.Y. revised the manuscript, contributed to the design of the experimental strategy, and provided overall support for the study; P.W.L.T. analyzed the data, drafted the manuscript, and prepared the figures; Q.W. designed constructs, drafted the manuscript and figures, and provided overarching support for the study.

---

**Table S1.** AAV productivity of AAVone systems in suspension cultured HEK 293T and HEK 293 cells.

| Cell lines | pAAVone plasmids | pAAVone size (kb) | AAV Serotype | AAV size (kb) | Scale (mL) | Cell density (1E6 cells/ml) | DNA Amount (ug/1E6 cells) | Transfection reagents (ratios with DNA) | Crude lysate titer (GC/L) | Purified vector yield (GC/L) |
| --- | --- | --- | --- | --- | --- | --- | --- | --- | --- | --- |
| HEK 293T | pAAVone-AAV1-CMV-EGFP-2.7kb | 15.7 | AAV1 | 2.7 | 130 | 3.0 | 0.75 | PEI<sub>Max</sub> (2:1) | 1.83E+15 | 2.84E+14 |
|  | pAAVone-AAV2-CMV-EGFP-2.7kb | 15.7 | AAV2 | 2.7 | 130 | 3.0 | 0.75 | PEI<sub>Max</sub> (2:1) | 4.90E+14 | 1.59E+14 |
|  |  |  |  |  | 160 | 3.0 | 0.5 | PEI<sub>Max</sub> (2:1) | 6.92E+14 | 1.46E+14 |
|  |  |  |  |  | 160 | 3.0 | 0.5 | PEI<sub>Max</sub> (3:1) | 7.91E+14 | 1.61E+14 |
|  |  |  |  |  | 400 | 3.0 | 0.5 | PEI<sub>Max</sub> (2:1) | 8.90E+14 | 2.14E+14 |
|  | pAAVone-AAV2-CMV-EGFP-2.2K | 15.2 | AAV2 | 2.2 | 90 | 3.0 | 0.5 | PEI<sub>Max</sub> (2:1) | / | 1.90E+14 |
|  | pAAVone-AAV5-CMV-EGFP-2.7kb | 15.7 | AAV5 | 2.7 | 130 | 3.0 | 0.75 | PEI<sub>Max</sub> (2:1) | 1.23E+15 | 1.65E+14 |
|  | pAAVone-AAV6-CMV-EGFP-2.7kb | 15.7 | AAV6 | 2.7 | 130 | 3.0 | 0.75 | PEI<sub>Max</sub> (2:1) | 2.01E+14 | 5.20E+13 |
|  | pAAVone-AAV8-CMV-EGFP-2.7kb | 15.7 | AAV8 | 2.7 | 130 | 3.0 | 0.75 | PEI<sub>Max</sub> (2:1) | 1.06E+15 | 1.66E+14 |
|  | pAAVone-AAV8-CMV-EGFP-2.2K | 15.2 | AAV8 | 2.2 | 90 | 3.0 | 0.75 | PEI<sub>Max</sub> (2:1) | 1.56E+15 | 3.29E+14 |
|  | pAAVone-AAV9-CMV-EGFP-2.7kb | 15.7 | AAV9 | 2.7 | 130 | 3.0 | 0.75 | PEI<sub>Max</sub> (2:1) | 3.70E+15 | 1.03E+15 |
|  |  |  |  |  | 2100 | 3.0 | 0.75 | PEI<sub>Max</sub> (2:1) | 2.04E+15 | 5.20E+14 |
|  |  |  |  |  | 90 | 3.0 | 1.0 | PEI<sub>Max</sub> (2:1) | 1.96E+15 | 4.77E+14 |
|  |  |  |  |  | 90 | 3.0 | 0.75 | PEI<sub>Max</sub> (2:1) | 2.49E+15 | 5.99E+14 |
|  |  |  |  |  | 90 | 3.0 | 0.5 | PEI<sub>Max</sub> (2:1) | 3.07E+15 | 6.80E+14 |
|  |  |  |  |  | 90 | 3.5 | 0.5 | PEI<sub>Max</sub> (2:1) | 3.01E+15 | 7.87E+14 |
|  |  |  |  |  | 90 | 4.0 | 0.5 | PEI<sub>Max</sub> (2:1) | 2.38E+15 | 4.22E+14 |
|  | pAAVone-AAV9-CMV-EGFP-4.4kb | 17.4 | AAV9 | 4.4 | 90 | 3.0 | 0.25 | PEI<sub>Max</sub> (2:1) | 2.26E+15 | / |
|  | pAAVone-AAV9-CMV-EGFP-4.6kb | 17.6 | AAV9 | 4.6 | 90 | 3.0 | 0.25 | PEI<sub>Max</sub> (2:1) | 1.76E+15 | / |
|  | pAAVone-AAV9-CMV-EGFP-4.7kb | 17.7 | AAV9 | 4.7 | 90 | 3.0 | 0.25 | PEI<sub>Max</sub> (2:1) | 1.94E+15 | / |
|  | pAAVone-AAVrh.10-CMV-EGFP-2.7kb | 15,719 | AAVrh.10 | 2.7 | 130 | 3.0 | 0.75 | PEI<sub>Max</sub> (2:1) | 1.61E+15 | 3.42E+14 |
| HEK 293 | pAAVone-AAV2-CMV-EGFP-2.2K | 15.2 | AAV2 | 2.2 | 30 | 3.0 | 0.35 | PEI<sub>Max</sub> (2:1) | 3.54E+14 | / |
|  | pAAVone-AAV6-CMV-EGFP-2.2K | 15.2 | AAV6 | 2.2 | 30 | 3.0 | 0.35 | PEI<sub>Max</sub> (2:1) | 2.91E+14 | / |
|  | pAAVone-AAV8-CMV-EGFP-2.2K | 15.2 | AAV8 | 2.2 | 30 | 3.0 | 0.35 | PEI<sub>Max</sub> (2:1) | 8.69E+14 | / |
|  | pAAVone-AAV9-CMV-EGFP-2.2K | 15.2 | AAV9 | 2.2 | 30 | 3.0 | 0.50 | PEI<sub>Max</sub> (2:1) | 1.55E+15 | / |
|  | pAAVone-AAV2-CMV-EGFP-2.2K | 15.2 | AAV2 | 2.2 | 30 | 3.0 | 0.375 | PEI<sub>Max</sub> (2:1) | 4.98E+14 | / |
|  |  |  |  |  | 30 | 3.0 | 0.25 | PEI<sub>Max</sub> (2:1) | 4.66E+14 | / |
|  |  |  |  |  | 30 | 3.0 | 0.375 | FectoVIR (1:1) | 4.42E+14 | / |
|  |  |  |  |  | 30 | 3.0 | 0.25 | FectoVIR (1:1) | 5.12E+14 | / |
|  |  |  |  |  | 30 | 3.0 | 0.375 | FectoVIR (2:1) | 6.48E+14 | / |
|  |  |  |  |  | 30 | 3.0 | 0.25 | PEI<sub>Max</sub> (2:1) | 6.53E+14 | / |
|  | pAAVone-AAV9-CMV-EGFP-2.2K | 15.2 | AAV9 | 2.2 | 30 | 3.0 | 0.35 | PEI<sub>Max</sub> (2:1) | 1.53E+15 | / |
|  |  |  |  |  | 30 | 3.0 | 0.35 | PEI<sub>Max</sub> (2:1) | 1.21E+15 | / |
|  |  |  |  |  | 30 | 3.0 | 0.50 | PEI<sub>Max</sub> (1:1) | 3.19E+15 | / |
|  |  |  |  |  | 30 | 3.0 | 0.30 | PEI<sub>Max</sub> (1:1) | 2.79E+15 | / |

**Table S2.** Summary of aligned SMRT reads to references.

| **References** | **pHelper**<br>**counts (%)** | **mini-pHelper**<br>**counts (%)** | **AAVone**<br>**counts (%)** |
| --- | --- | --- | --- |
| GOI from ITR-to-ITR | 126,756 (96.04%) | 115,005 (96.62%) | 190,658 (96.90%) |
| GOI Full Reference | 127,018 (96.23%) | 115,164 (96.75%) | 191,168 (97.16%) |
| GOI from ITR-to-ITR Carrying Backbone | 5,209 (3.95%) | 3,204 (2.69%) | 1,715 (0.87%) |
| GOI backbone Only | 152 (0.12%) | 85 (0.07%) | 105 (0.05%) |
| Rep Only | 15 (0.01%) | 3 (0.00%) | 10 (0.01%) |
| Rep Chimeric with GOI | 377 (0.29%) | 243 (0.20%) | 89 (0.05%) |
| Cap only | 28 (0.02%) | 28 (0.02%) | 29 (0.01%) |
| Cap Chimeric with GOI | 336 (0.25%) | 240 (0.20%) | 201 (0.10%) |
| Rep and Cap overlap | 51 (0.04%) | 18 (0.02%) | 66 (0.03%) |
| Rep-Cap Carrying Backbone | 63 (0.05%) | 51 (0.04%) | N/A |
| Rep-Cap Backbone | 224 (0.17%) | 14 (0.01%) | N/A |
| Rep-Cap Carrying with GOI | 687 (0.52%) | 466 (0.39%) | 319 (0.16%) |
| E2 only | 63 (0.05%) | 34 (0.03%) | 90 (0.05%) |
| E2 chimeric with GOI | 389 (0.29%) | 335 (0.28%) | 610 (0.31%) |
| E4 only | 12 (0.01%) | 11 (0.01%) | 34 (0.02%) |
| E4 chimeric with GOI | 147 (0.11%) | 150 (0.13%) | 260 (0.13%) |
| VA RNA only | 1 (0.00%) | 0 (0.00%) | 0 (0.00%) |
| VA RNA chimeric with GOI | 131 (0.10%) | 123 (0.10%) | 4,614 (2.35%) |
| pHelper/mini-pHelper carrying backbone | 278 (0.21%) | 181 (0.15%) | N/A |
| pHelper/mini-pHelper backbone | 0 (0.00%) | 0 (0.00%) | N/A |
| Backbone combined(AAVone only) | N/A | N/A | 7,950 (4.04%) |
| hg38 | 4,453 (3.37%) | 3,471 (2.92%) | 5,311 (2.70%) |
| hg38 chimeric with GOI | 677 (0.51%) | 527 (0.44%) | 814 (0.41%) |
| unmapped reads | 921 (0.70%) | 715 (0.60%) | 990 (0.50%) |
| Total non-AAV containing reads | 4,697 (3.56%) | 3,659 (3.07%) | 5,487 (2.79%) |
| Total plasmid impurities | 8,303 (6.29%) | 5,591 (4.70%) | 7,899 (4.01%) |
| Total reads | 131,989 | 119,029 | 196,753 |

**Figure S1.**
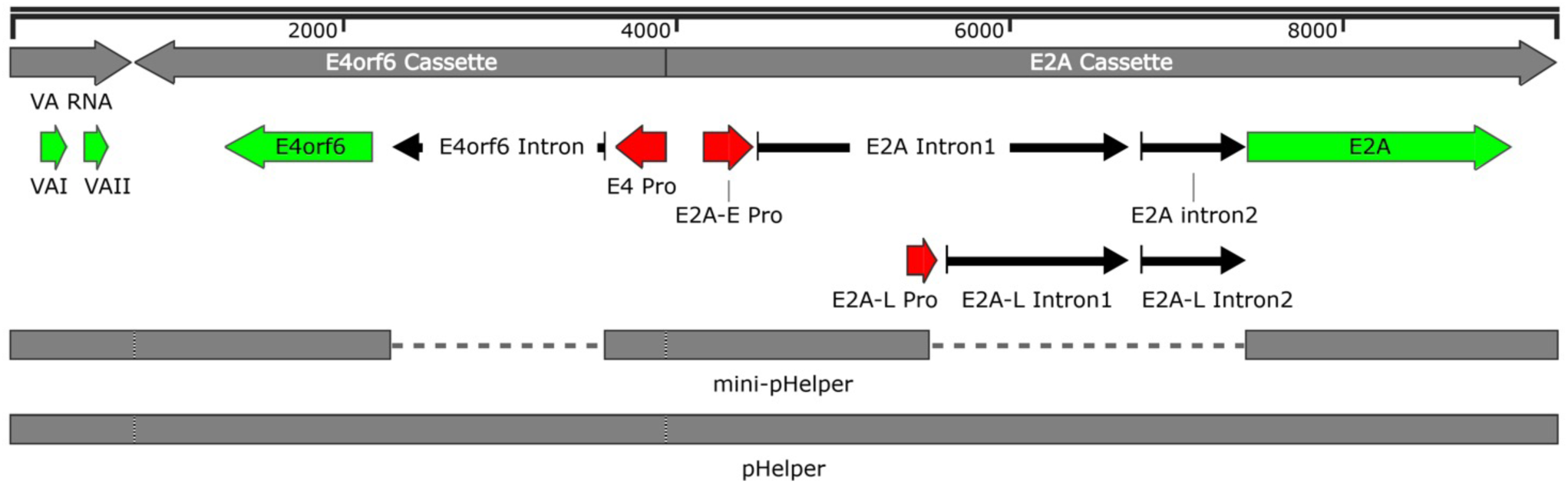
Construction of mini-pHelper. The original pHelper contains 9.3 kb of the AdV2 genome with three expression cassettes (gray), including *VA RNA* (731 bp), *E4orf6* (3,192 bp) and *E2A* (5,343 bp). The *E2A* and *E4orf6* components are the main contributors to the plasmid’s large size. The *E2A* gene transcripts: early (E2A-E) and late (E2A-L) mRNAs share a small second intron, but have different first intron. The *E4orf6* has a 1,275 bp intron. In the mini-pHelper, intron 1 and 2 of E2A-L, as well as the intron of *E4orf6* were eliminated, resulting in a 8.4-kb plasmid with 6.1 kb of AdV2. Red, promoters; green, ORFs; arrows, introns.

**Figure S2.**
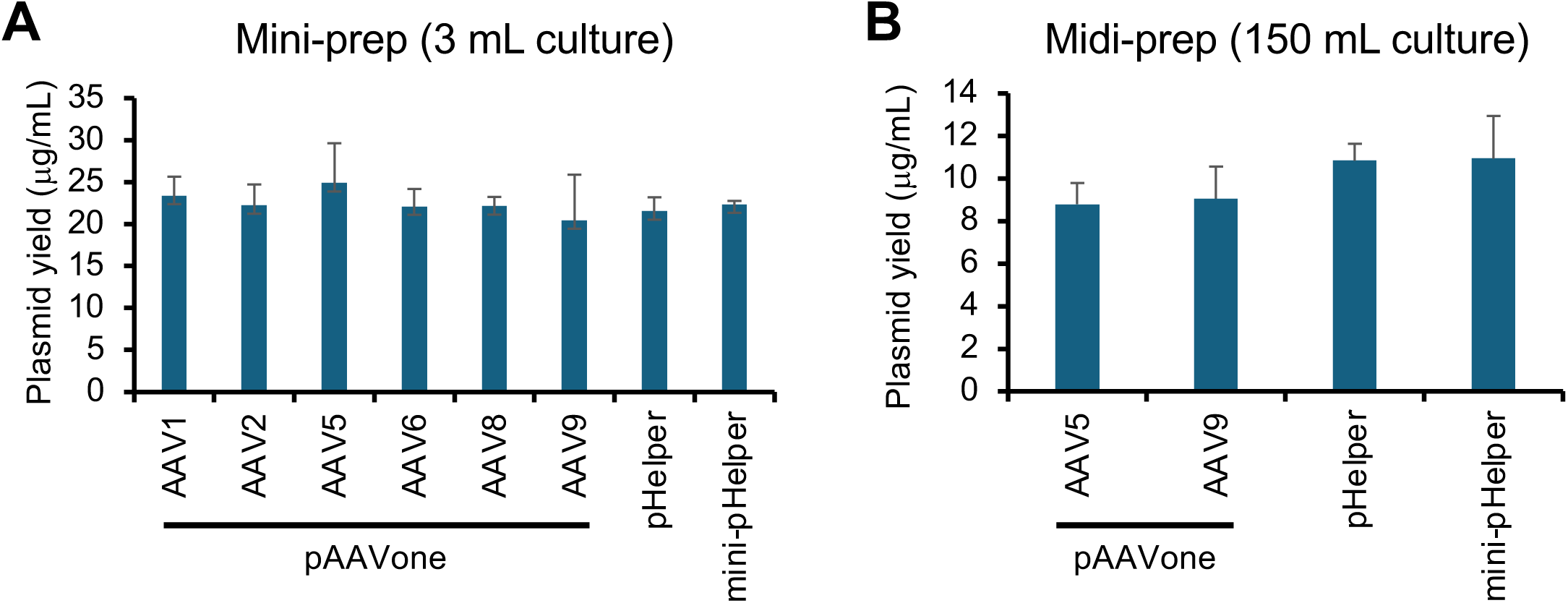
Examination of pAAVone plasmid yields. Comparisons of plasmid concentrations produced by mini-preps **(A)** and midi-preps **(B)** across multiple serotypes.

**Figure S3.**
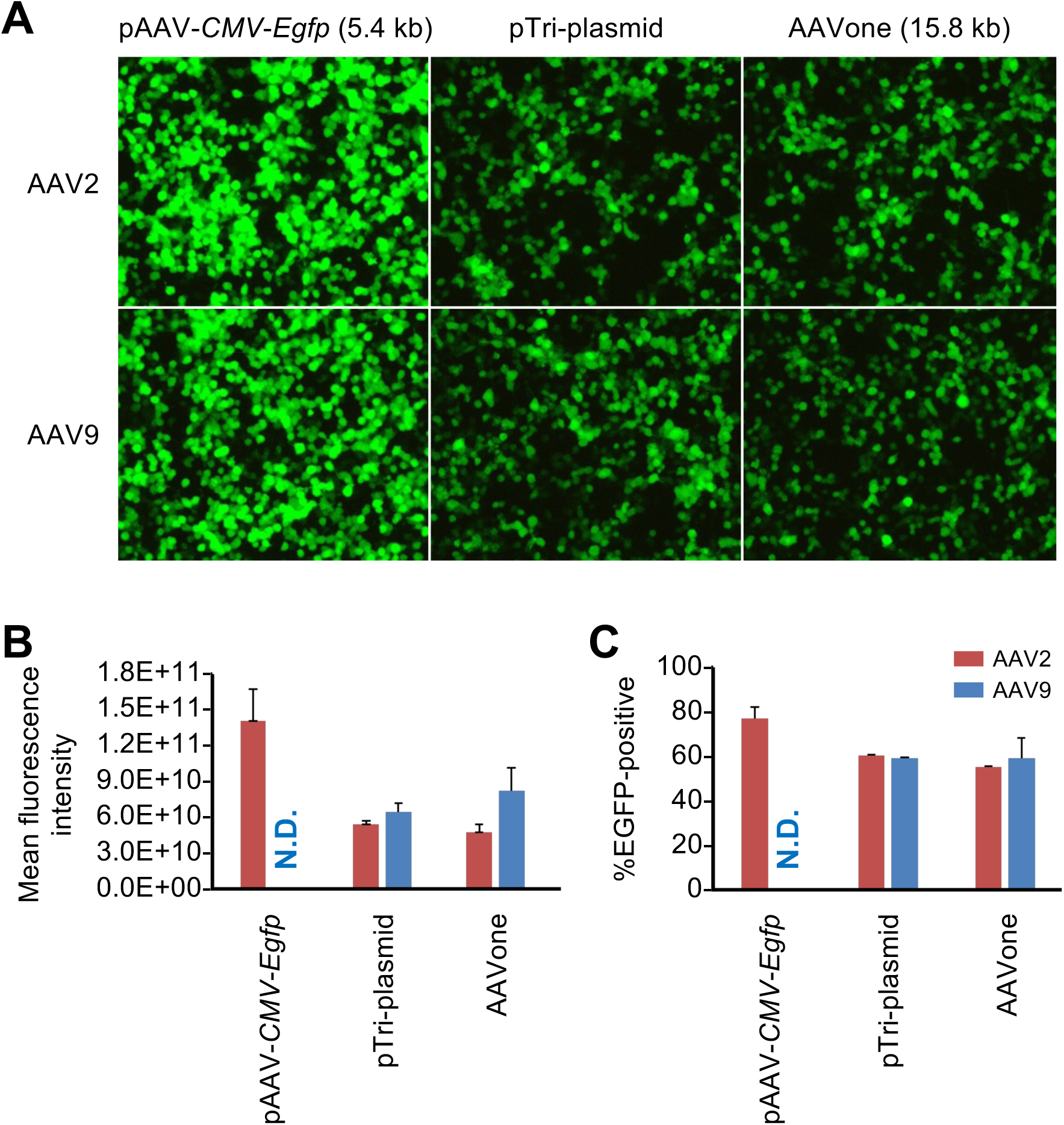
Transfection efficiency of the pAAV-CMV-Egfp, pTri-plasmid system and the pAAVone in adherent HEK 293T cells. At 48-h post-transfection, EGFP was observed by epifluorescence **(A),** and the mean fluorescence intensity **(B)** and percentage of EGFP-positive cells **(C)** were analysis with flow cytometry.

**Figure S4.**
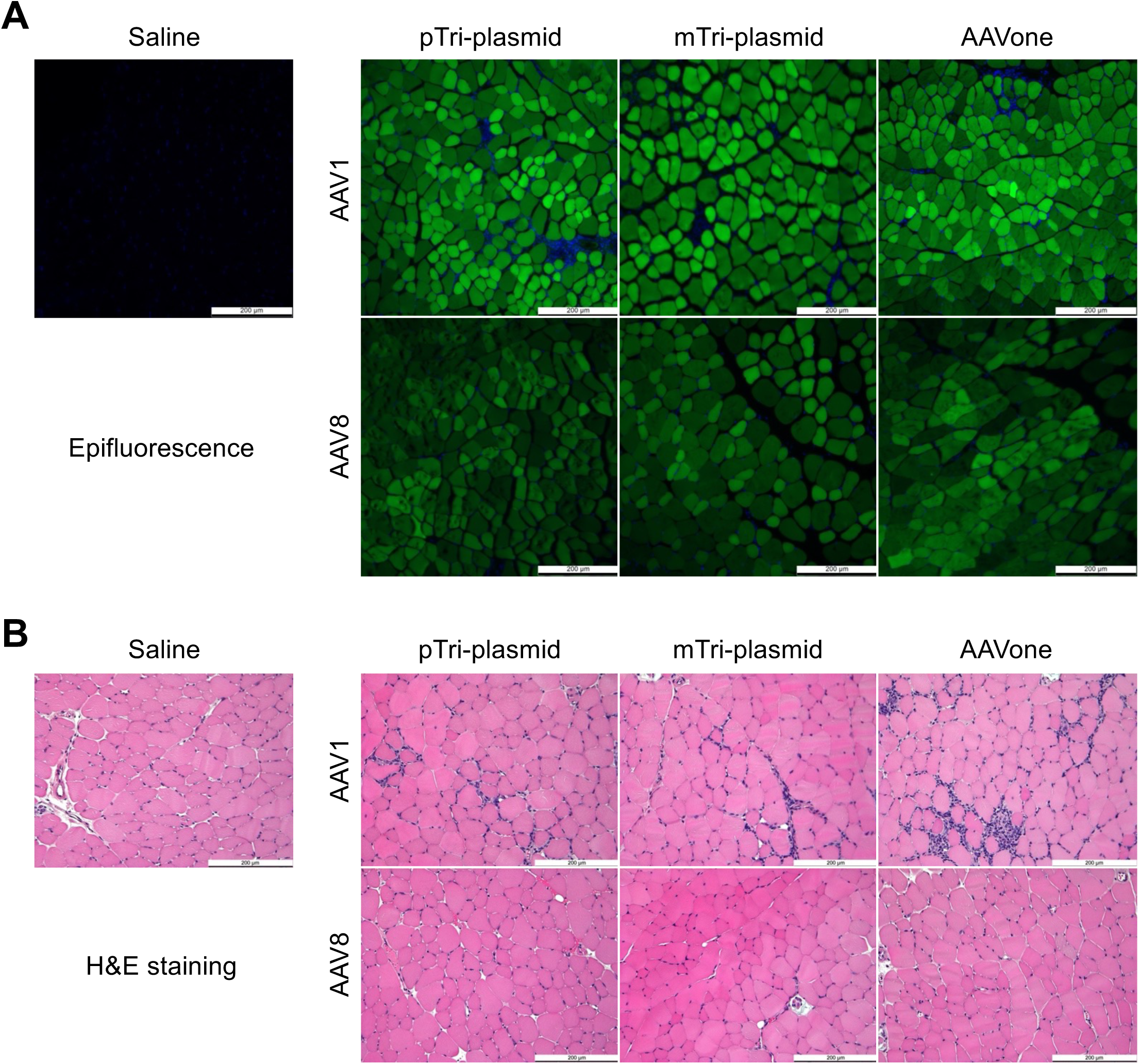
AAVone-packaged AAV2 and AAV8 vectors exhibit similar muscle transduction efficacies to vectors packaged by triple-plasmid transfection. Representative epifluorescence microscopy images **(A)** and H&E staining **(B)** of TA cross-sections at 2 weeks post-injection with ssAAV1.*CMV.Egfp* (top row) and ssAAV8.*CMV.Egfp* (bottom row) produced via the three three packaging systems. EGFP (green) and nuclei (blue). Tissues from mice injected by saline solution serve as negative controls.

**Figure S5.**
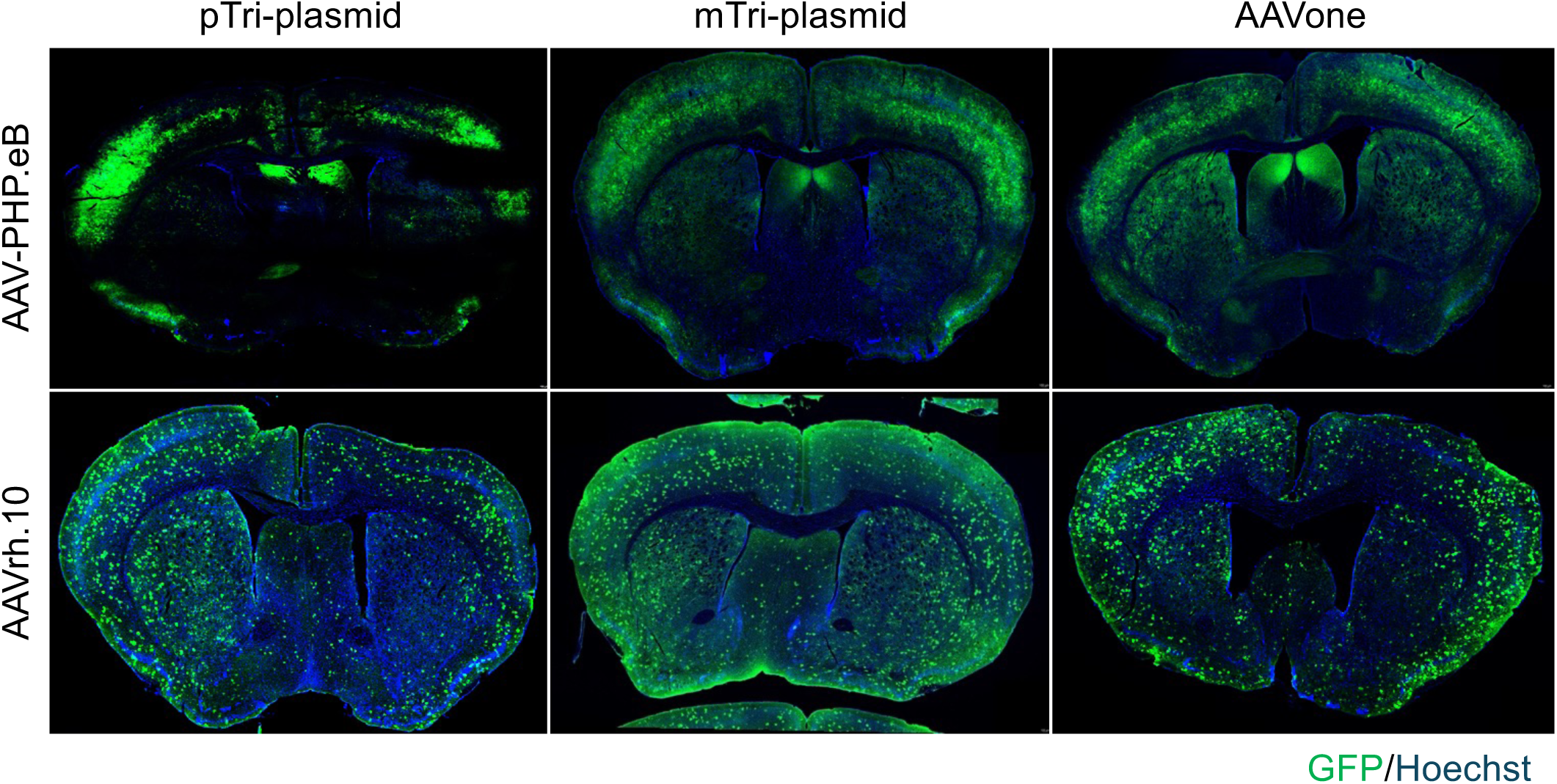
AAVone-packaged AAV-PHP.eB and AAVrh.10 vectors exhibit similar brain transduction efficacies to vectors packaged by triple-plasmid transfection. Epifluorescence images of mouse brain cryosections treated with AAV-PHP.eB vectors (top row) or AAVrh.10 vectors (bottom row) produced by AAVone or the triple-plasmid systems. Sections were stained with anti-GFP antibodies and Hoechst dye to label nuclei.

**Figure S6.**
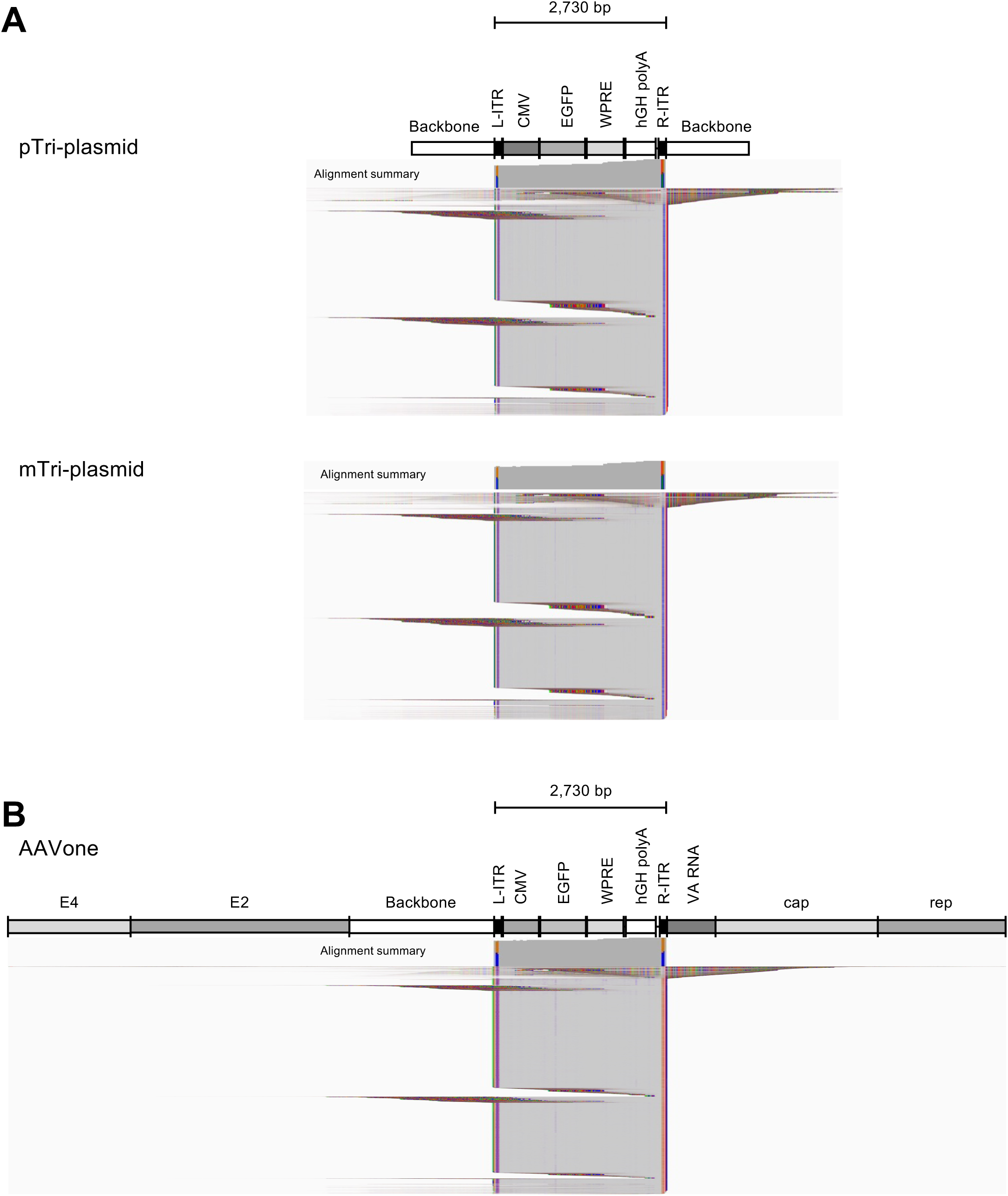
Genome integrity analysis of vectors produced by tri-plasmid systems and AAVone. **(A, B)** IGV display of SMRT reads representing AAV vector genomes aligned to the standard *cis* plasmid **(A)** from the pTri-plasmid (top) and the mTri-plasmid (bottom) packaging systems, and the AAVone **(B)** reference genomes. Both constructs carry the *CMV-Egfp* cassette. Reads are shown in squished displays with soft-clipped bases visible. Read matches (gray), mismatches (colored), and insertions/deletions (speckles) are shown. Alignment coverages are shown above each display.

**Figure S7.**
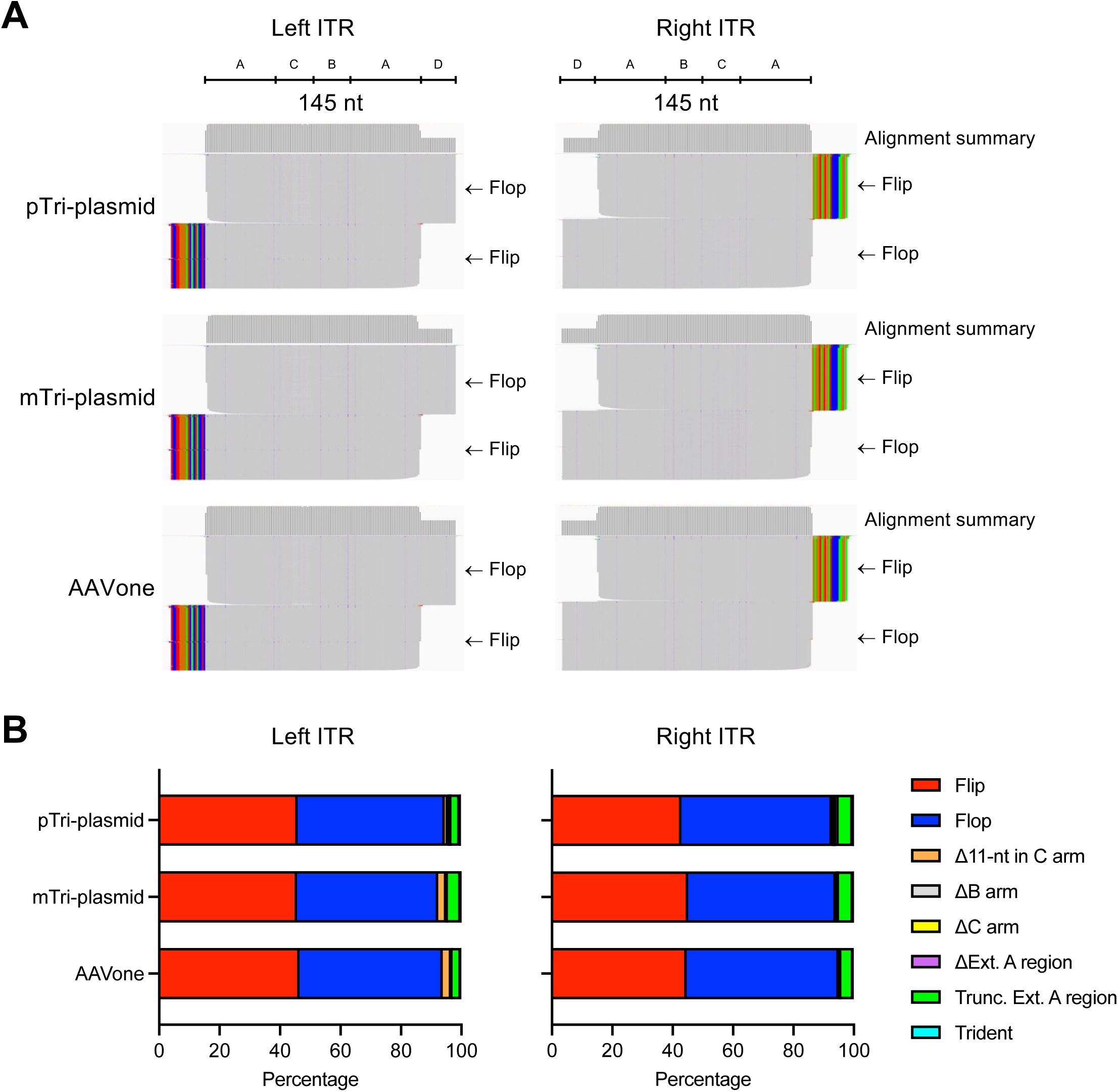
Repair of the Δ11-nt C arm ITR. **(A)** IGV display of left and right ITR regions extracted from SMRT reads from pTri-plasmid-, mTri-plasmid-, and AAVone-produced vectors aligned to the 145-nt flop-oriented ITR references. Alignments are shown in squished displays with soft-clipped bases visible. Read matches (gray), mismatches (colored), and insertions/deletions (speckles) are shown. Alignment coverages are shown above each display. **(B)** Distributions of ITR configurations at the left and right ITRs from the three production platforms. Flip (red); Flop (blue); 11-nt deletion, Δ11-nt in C arm (orange); B arm deletion, ΔB arm (grey); C arm deletion, ΔC arm (yellow); external A region deletion, ΔExt. A region (purple); truncation of the A region, Trunc. Ext. A region (green); and trident-shaped ITR (cyan).

**Figure S8.**
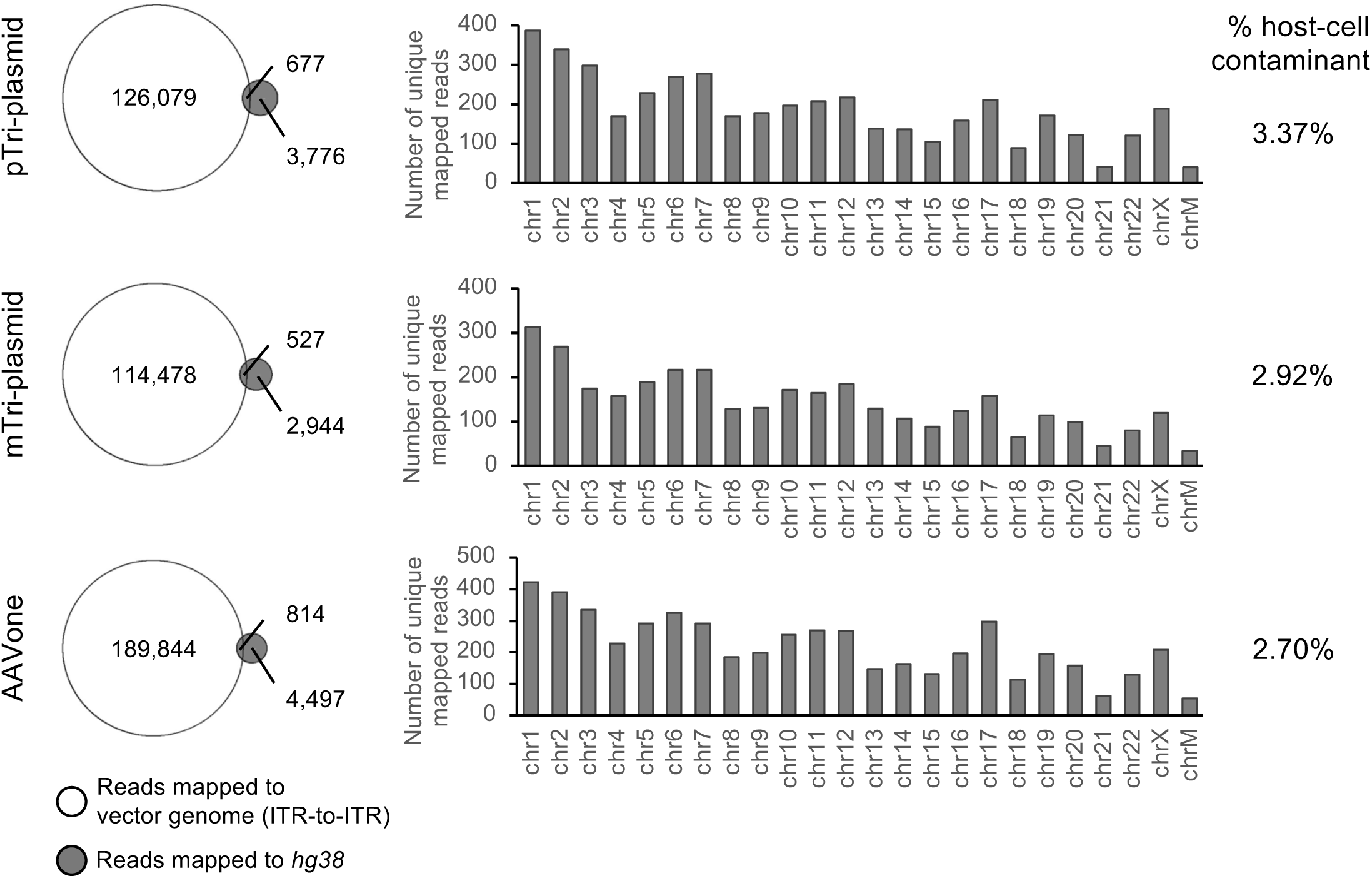
Profiling of hcDNA contaminants associated with triple-plasmid systems and AAVone. The Venn diagrams (left) show mapped reads to vector genome and to human genome hg38, as well as chimeric mapped reads for the pTri-plasmid (top), mTri-plasmid (middle), and AAVone (bottom) packaging systems. The number of reads mapped exclusively to the vector genome (ITR-to-ITR, white) and hg38 (human genome) are shown. The distribution of mapped reads to individual chromosomes can also be observed in the bar graph.

**Figure S9.**
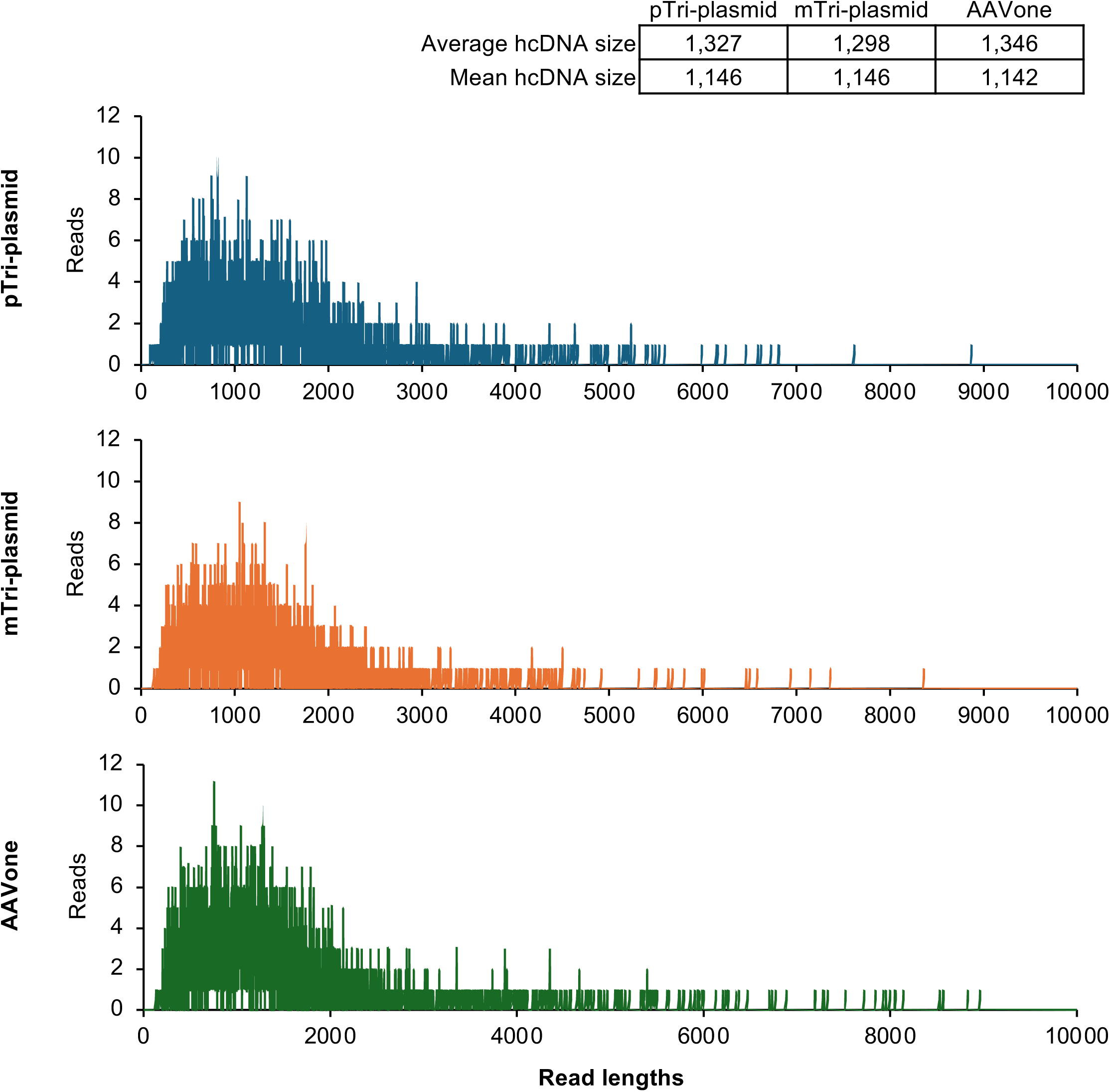
Length distributions of SMRT reads that map to hg38. Reads from vectors produced by the pTri-plasmid (top), mTri-plasmid (middle), and AAVone (bottom) systems are shown. A table summarizing the average amdn mean sizes are shown.

**Figure S10.**
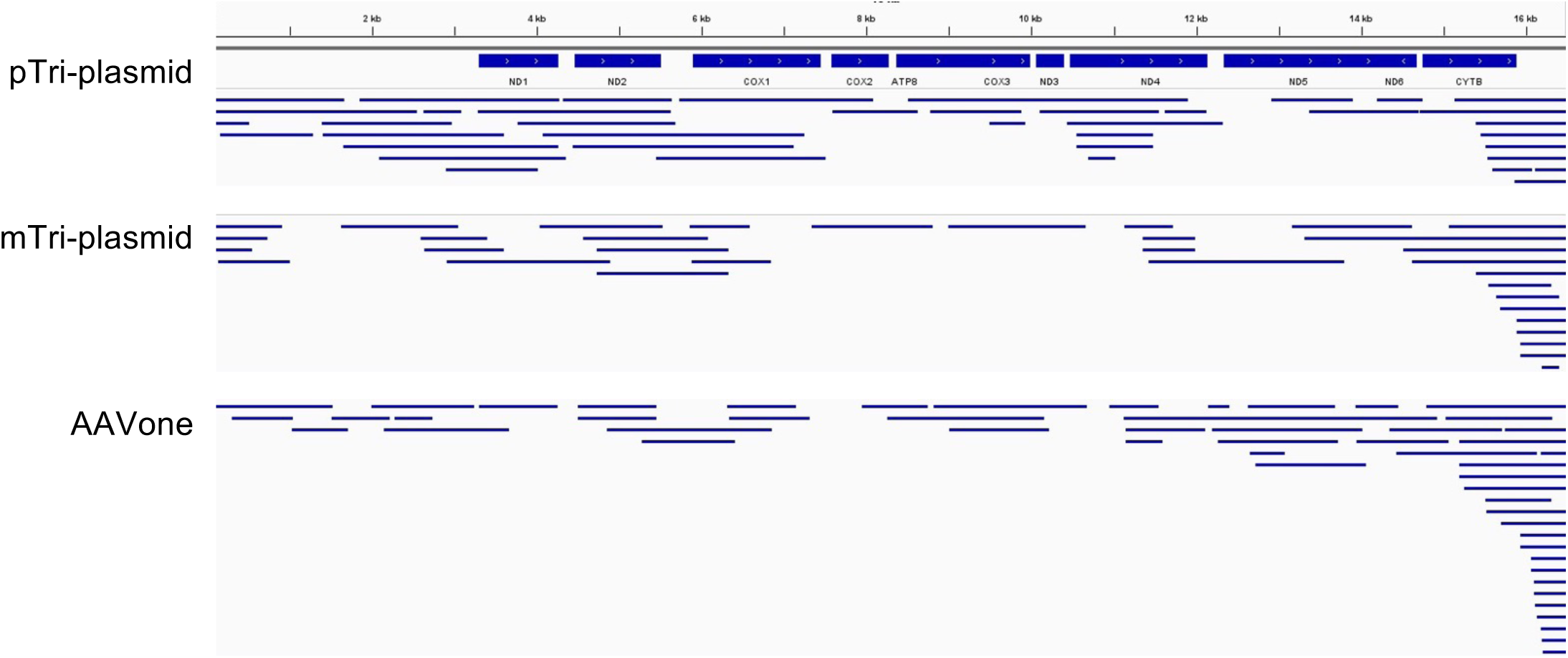
Alignment of SMRT reads from DNase-resistant DNA isolated from vectors to chrM. Reads from vectors produced by the pTri-plasmid (top), mTri-plasmid (middle), and AAVone (bottom) systems are shown mapping to the mitochondrial genome. Tracks of the mitochondrial genes are shown above.

**Figure S11.**
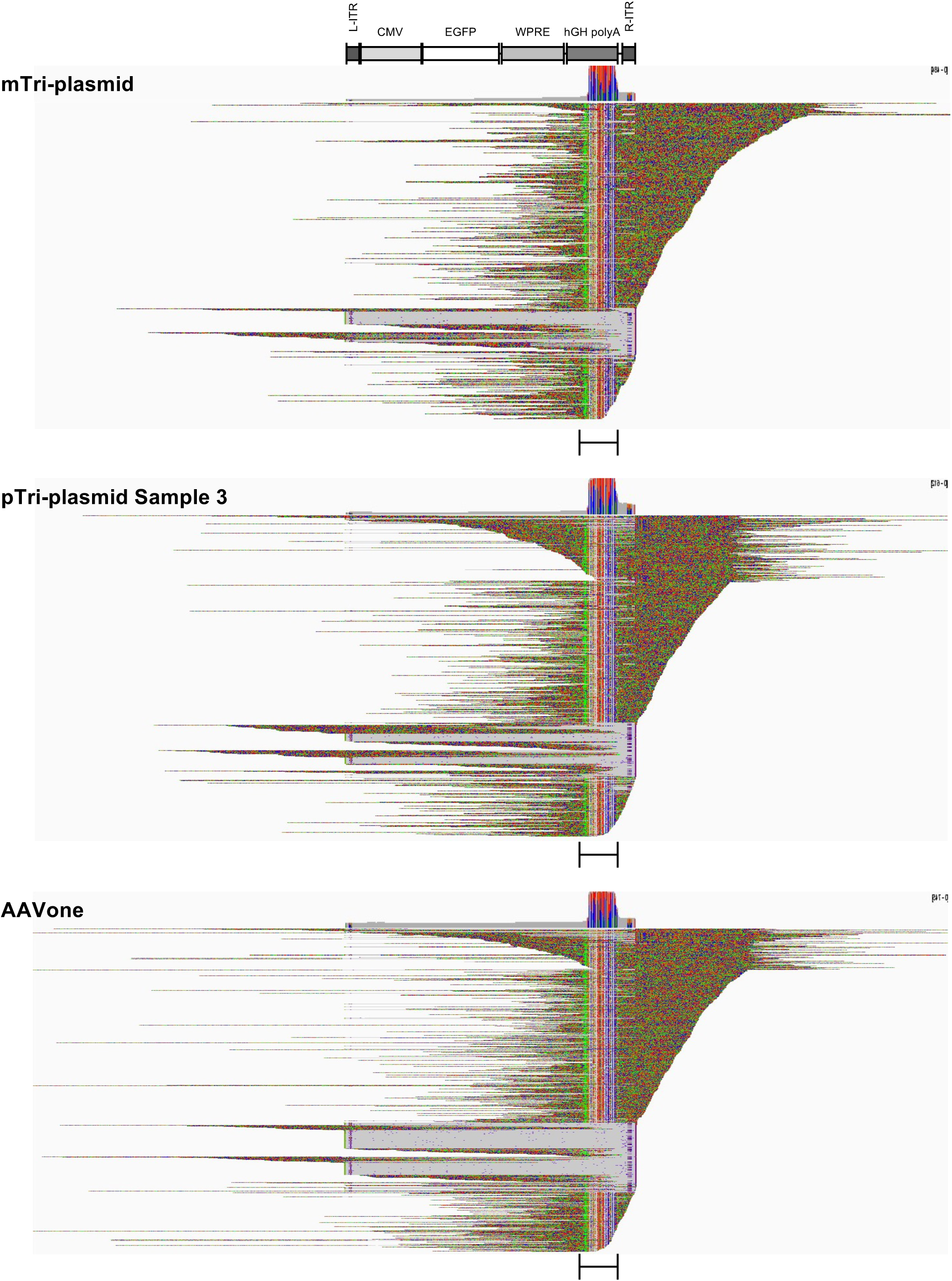
Alignment of chimeric reads to the GOI reference. Reads from vectors produced by the pTri-plasmid (top), mTri-plasmid (middle), and AAVone (bottom) systems are shown. Alignments are shown in squished displays with soft-clipped bases visible. Read matches (gray), mismatches (colored), and insertions/deletions (speckles) are shown. Multiple **r**eads falsely map to the human GH polyA (bracket).

**Figure S12.**
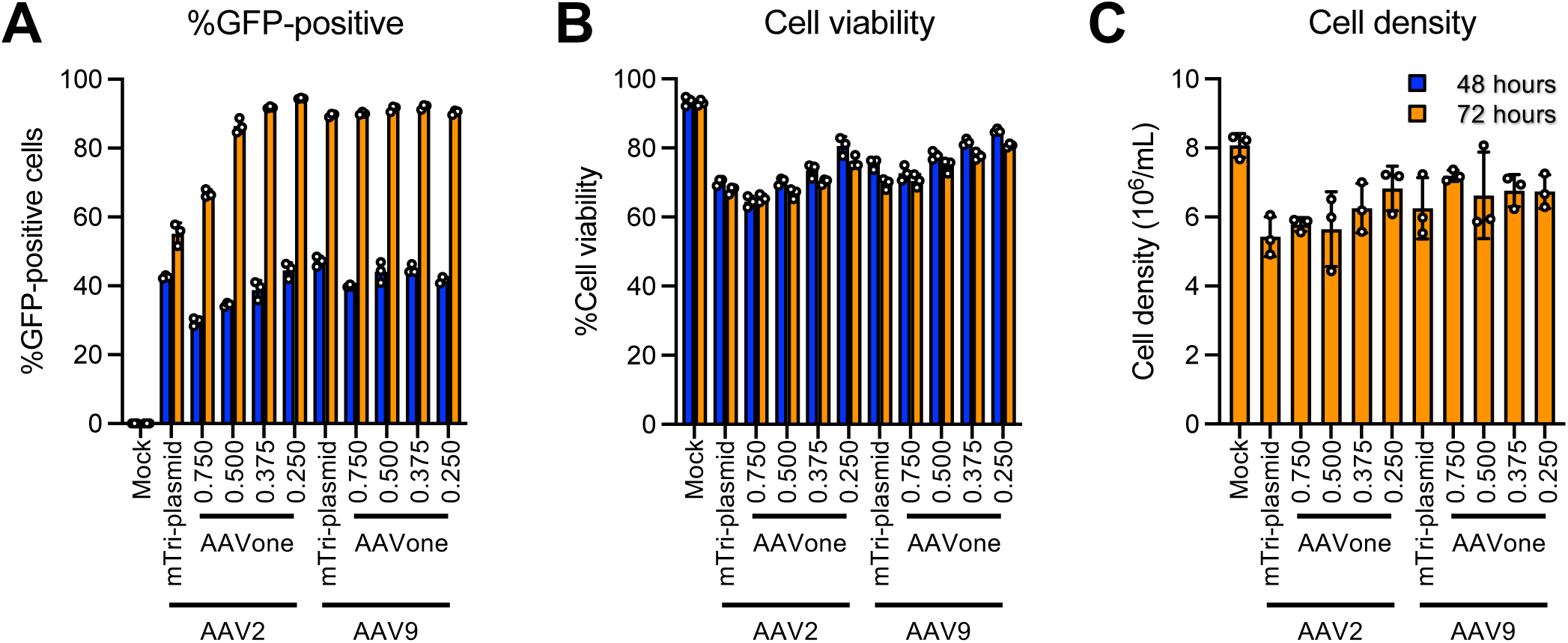
Reduction of AAVone plasmid used in production increases vector titers. **(A-C)** Cell metrics following transfection with different amounts of packaging plasmids. Percentage of EGFP-positive cells **(A),** cell viability **(B),** and cell density **(C)** are reported at 48-hours (blue bars) or 72-hours (orange bars) post-transfection.

**Figure S13.**
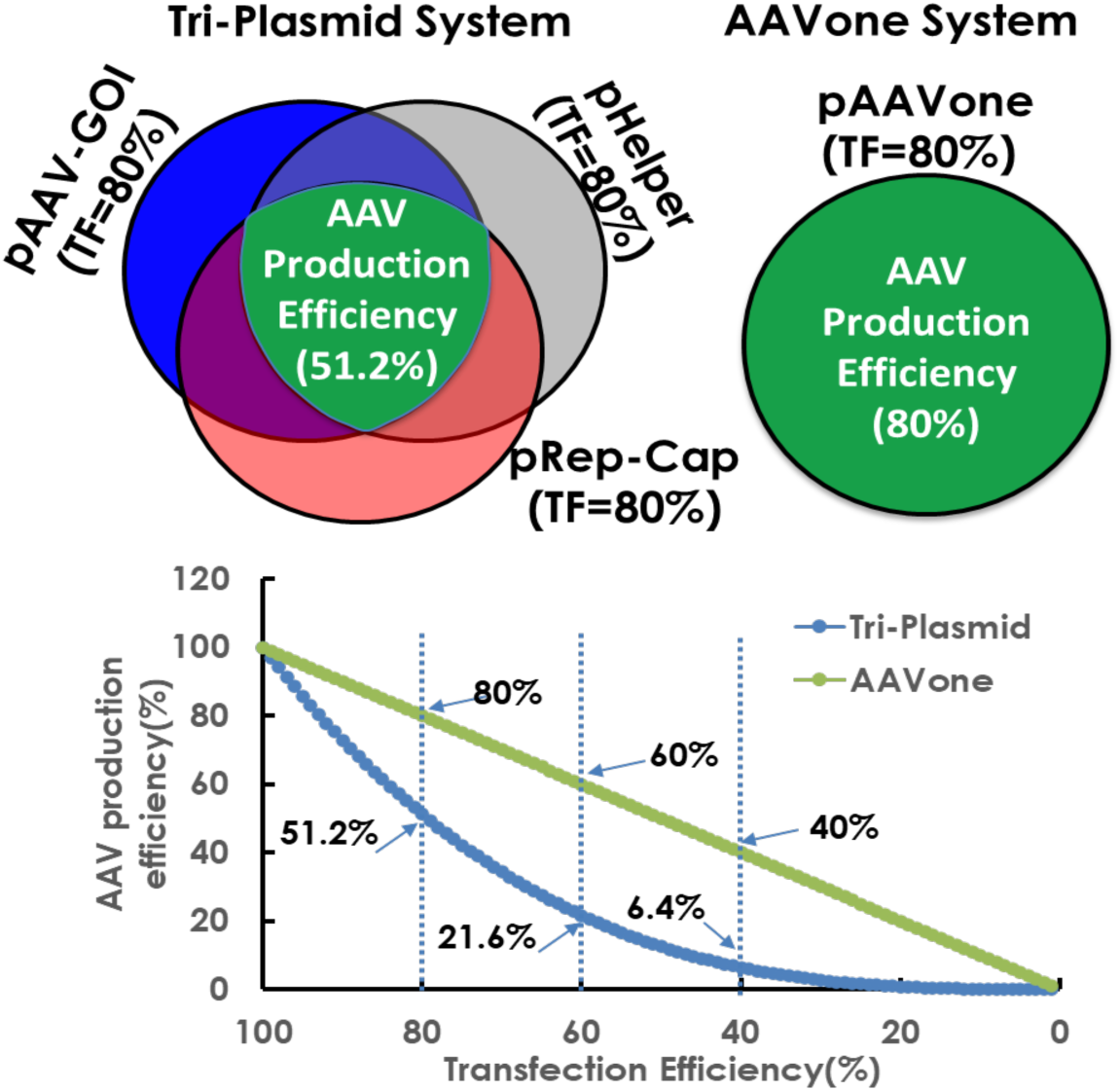
Theoretical relationship of transfection (TF) efficiency and AAV production efficiency with AAVone and triple-plasmid systems.

